# The mRNA covalent modification dihydrouridine regulates transcript turnover and photosynthetic capacity during plant abiotic stress

**DOI:** 10.1101/2025.01.17.633510

**Authors:** Li’ang Yu, Giovanni Melandri, Anna C. Nelson Dittrich, Sebastian Calleja, Diep R. Ganguly, Kyle Palos, Emily K. Brewer, Hillary Fischer, Bruno Rozzi, Aparna Srinivasan, Toshihiro Obata, Hamada Abd Elgawad, Gerrit T.S. Beemster, Riley Henderson, Ciara Denise Garcia, Xiaodan Zhang, David Stern, Andrea Eveland, Eric Lyons, A. Elizabeth Arnold, Aleksandra Skirycz, Susan J. Schroeder, Brian D. Gregory, Duke Pauli, Andrew D. L. Nelson

## Abstract

RNA Covalent Modifications (RCMs) are post-transcriptional chemical alterations that influence RNA stability and translation efficiency, thus play critical roles in eukaryotic growth and development. However, their role in regulating plant performance under abiotic stress remain largely unexplored. Here, we integrated multi-omics data in six *Sorghum bicolor* accessions under water-limiting conditions in the field to explore the relationship between RCMs and drought response. Within a stress and photosynthesis-associated gene co-expression module, we identified *SbDUS2*, a member of family of enzymes, conserved across eukaryotes, which catalyzes the reduction of uracil to dihydrouridine (DHU) on RNA molecules. DHU-modified transcripts in this module were enriched for photosynthetic functions and showed strong correlation with photosynthetic traits. To elucidate the function of this RCM, we characterized loss of function *dus2* mutants in the genetic model, *Arabidopsis thaliana*. Under control conditions, these DHU-deficient mutants exhibited impaired germination and delayed development. Furthermore, when exposed to heat or water-limiting conditions, these mutants showed significantly reduced net CO_2_ assimilation and survival. Using multiple transcriptome-wide RNA stability assays, we demonstrated that transcripts associated with lower DHU level in a *dus2* background generally exhibited increased stability compared to Col-0 controls. Particularly, lack of DUS2 led to the hyperstability of photosynthesis-related transcripts, impeding their turnover and likely preventing proper photosynthetic acclimation during stress. We propose a model based on these data where DHU acts as a critical post-transcriptional regulator marking mRNAs for rapid turnover under stress, highlighting an overlooked regulatory layer contributing to plant resilience.

## Introduction

Agricultural production faces intensifying challenges as drought and heat stressors increase in frequency and severity across major cropping regions (Alizadeh et al., 2020). These abiotic stressors trigger a range of adverse phenotypic and physiological responses, including heightened susceptibility to biotic stresses, altered development, and reduced yield. Consequently, enhancing plant productivity and resilience under such conditions has become a primary goal in improving stress tolerance. A widely adopted strategy to achieve this goal involves the exploration and utilization of genetic variation (Glaszmann et al., 2010; Khoury et al., 2022). While large population-level studies within many crop species have identified strong abiotic stress tolerance phenotypes and associated loci useful for breeding and selection (Gilliham et al., 2017), the crosstalk between many upstream regulatory genes and downstream trait-associated genes remains unclear and largely unexplored. In addition to feedback at the DNA, RNA, and protein level, there are additional levels of regulatory control that make identifying beneficial alleles for crop improvement more challenging.

RNAs endure numerous co- and post-transcriptional events that influence their function, one of which is the covalent modification, or isomerization, of RNA bases. RNA covalent modifications (RCMs) are an emerging and underexplored layer of regulation in eukaryotes (Harcourt et al., 2017; Roundtree et al., 2017). RCMs are widespread and chemically diverse, decorating all classes of RNA, non-coding and coding alike (Finet et al., 2022a). As a natural result of their higher abundance, most RCMs were first characterized on housekeeping RNAs such as tRNAs and rRNAs (Yu et al., 2011; Toubdji et al., 2024). One such RCM is dihydrouridine (DHU), which is formed by the reduction of the double bond in uridine by a highly conserved class of dihydrouridine synthase (DUS) enzymes (Yu et al., 2011). DHU is known to disrupt base stacking and increase RNA flexibility, particularly in tRNAs, where it is critical for the formation of the D-loop (Toubdji et al., 2024), a role that is conserved across all living organisms (Lombard et al., 2022). In non-plant systems, DHU has recently been found in mRNAs, predominantly in transcripts associated with core cellular mechanisms (Kato et al., 2005; Rider et al., 2009; Finet et al., 2022b). Disruption or depletion of DHU from these transcripts results in growth defects in yeast and, in humans, enhanced tumorigenesis (Kato et al., 2005; Rider et al., 2009). Despite the growing evidence that DHU plays an important role in mRNAs, how DHU regulates mRNA function, and the extent to which this RCM is present in plant mRNAs, is unclear.

Much of what we know about mRNA-associated RCMs comes from studies on N6-methyladenosine (m^6^A), which has been shown to influence plant development and stress responses by modulating mRNA fate (Zhou et al., 2022; Shao et al., 2025; Liu et al., 2025). RCMs are known to regulate RNA stability, structure, translation, and localization within, and potentially between, cells (Arribas-Hernández et al., 2018; Anderson et al., 2018; Luo et al., 2020; Cheng et al., 2021; Prall et al., 2023). Despite this growing appreciation of the roles that RCMs can perform, many evolutionarily conserved RCMs, such as DHU, pseudouridine (Ψ), N^1^-methyladenosine (m^1^A), and 5-methylcytidine (m^5^C), remain largely unknown in plants (Tang et al., 2020; Yang et al., 2020; Li et al., 2025). This knowledge gap stems in part from technical barriers. Several RCMs can be detected with antibody-based methods (Dominissini et al., 2012; Meyer et al., 2012; Edelheit et al., 2013; Li et al., 2016), and more recently at the transcript level using Oxford Nanopore direct RNA-seq (DRS) along with trained algorithms to detect specific modified ribonucleotides (Garalde et al., 2018; Liu et al., 2019; LaPierre et al., 2019; Wu et al., 2024; Cruciani et al., 2025). For modifications that impact the base-pairing edge, bioinformatic approaches that interpret the pattern of resulting RT-induced errors at modified sites have been deployed with great success (Ryvkin et al., 2013; Tan et al., 2021). Although somewhat limited in resolution (Helm and Motorin, 2017), these approaches have nevertheless been helpful in extending insights into the roles of less studied RCMs (Sharma et al., 2023).

Sorghum [*Sorghum bicolor* (L.) Moench] is a heat- and drought-tolerant crop widely cultivated for grain, forage, and cellulosic biomass production (Lobell et al., 2008; Abreha et al., 2021). Due to its phylogenetic placement within the *Poaceae*, findings from sorghum often transfer to other critical agronomic crops such as *Zea mays*. A well annotated genome (McCormick et al., 2018), along with the physiological and morphological variation found within multiple diversity panels (Morris et al., 2013; Wu et al., 2022; Boatwright et al., 2022), make sorghum ideal for supporting comparative, systems-level analyses on field-grown plants. Indeed, several studies have used these resources to highlight transcriptional responses to heat and drought stress in sorghum, uncovering genetic factors that might contribute to improved stress resilience in sorghum and other important grasses (Boyles et al., 2019; Varoquaux et al., 2019). Thus, these resources and characteristics make sorghum ideally suited for uncovering RNA-based regulatory mechanisms contributing to plant resilience.

Here we leveraged a systems biology approach to study how six phylogenetically distinct sorghum accessions with different levels of drought tolerance respond to water limiting conditions over time. Our integrative approach using global-wide datasets identified the transcripts with DHU modification and its corresponding enzyme for DHU synthesis in sorghum, SbDUS2, as strongly associated with plant photosynthetic performance in the field. Interestingly, SbDUS2 is orthologous to human and fission yeast DUS2, which were both previously found to be responsible for mRNA DHU deposition. Given the deeply conserved nature of the DUS enzymes, we turned to *Arabidopsis thaliana*, where genetic *loss-of-function* mutants for the orthologous DUS2 were available, to investigate the functional role of DHU in *Arabidopsis*. Under control conditions, *Atdus2* mutants displayed reduced germination and delayed growth relative to *AtDus2* controls. A set of complementary approaches were used to confirm that Arabidopsis *dus2* mutants displayed reduced DHU levels, particularly on mRNAs associated with photosynthesis and metabolism. After confirming that *Atdus2* mutants displayed reduced DHU levels on mRNAs, we used this background to better understand the role of DHU in plant stress responses. Interestingly, *Atdus2* mutants exhibited a similar physiological response to drought and heat, showing reduced CO₂ assimilation and lower survival. In Col-0 controls under heat, but not in the *Atdus2* background, photosynthesis-related transcripts were found to be differentially DHU modified, a phenomenon that coincided with decreased abundance for the associated transcripts. Using transcriptional arrest time course-derived RNA decay assay at global level, we demonstrated that DHU predominantly destabilizes mRNAs under control and abiotic stress conditions in Arabidopsis, and propose a model whereby DHU is critical for mRNA turnover, particularly under stress scenarios, highlighting the significance of this overlooked regulatory mechanisms with real-world agronomic relevance.

## Methods and Materials

### Sorghum growth and phenotypic measurement

Six selected sorghum accessions (PI 533871: M1, PI 533961: DL/59/1530, PI 656041: 80M, PI 656053: N290-B, PI 656076: SC1271, and PI 656096: SC391) were grown at the University of Arizona Maricopa Agricultural Center (MAC). All plants were well-watered (WW) at the same rate (∼24% soil volume metric water content, SVWC), corresponding approximately to the soil field capacity) until flag leaf appearance in the whorl (∼47 days after planting), after which irrigation was reduced to a subset of plots to mimic drought treatment (water-limited, WL; 16% SVWC). The destructive plant phenotyping was conducted at the end of the season (maturity stage). Measured traits included above-ground biomass, root biomass, and three-leaf weights. For other time-series molecular phenotypes, samples were collected weekly (on Thursdays) for seven weeks (week 1-week 7, from August 13^th^ to September 24^th^, 2020) on a specific range of a large field experiment conducted at the University of Arizona’s Maricopa Agricultural Center (MAC). For transcriptomics, metabolomics, and analysis of other biochemical traits, samples consisting of six leaf disks from five different plants per accession, treatment, and time point were collected (from 10:30 to 11:30) in three different 1.5 mL tubes and immediately snap-frozen in liquid nitrogen. LI-6800 derived photosynthesis-related traits (*N =* 29, LI-COR Lincoln, NE) were measured in the morning (from 9:00 to 10:00) and afternoon (from 13:00 to 14:00) with the following settings: Flow rate = 600 µmol s^-1^; Mixing fan speed =12,000 rpm; CO_2_ = 420 ppm; Light= 2,000 µmol m^-2^ s^-1^. The youngest fully expanded leaf on the main stem of each plant was selected for the measurements and data were logged as soon as assimilation and stomatal conductance curves (monitored in real-time on the instrument display) when reaching a plateau (on average 30 seconds after a leaf was clamped).

### Sorghum biochemical trait measurements

Highly concentrated metabolites (e.g., total soluble sugars), levels of oxidative stress markers and molecular antioxidants, and antioxidant enzyme and photorespiration enzyme activities were analyzed using existing protocol (Melandri et al., 2020). Chlorophyll and carotenoid contents were measured after acetone extraction (Porra et al., 1989). Sugar levels were quantified using enzyme-coupled assays (Riewe et al., 2012). Oxidative stress markers, such as malondialdehyde (MDA) and protein carbonyl content, were measured to assess lipid peroxidation and protein oxidation (Levine et al., 1994; Hodges et al., 1999). Photorespiration-related enzyme activities, glycolate oxidase (GOX) and hydroxypyruvate reductase (HPR), were assessed based on established methodologies (Feierabend and Beevers, 1972; Schwitzguebel and Siegenthaler, 1984). Lipoxygenase (LOX) was extracted in potassium phosphate buffer (pH 7.0), and its activity was measured by changes in conjugated dienes. Total antioxidant capacity (TAC), total polyphenol content (Poly), and flavonoids were evaluated using standard spectrophotometric assays (Benzie and Strain, 1999; Zhang et al., 2006). Enzyme activities of key antioxidants, including ascorbate peroxidase (APX), dehydroascorbate reductase (DHAR), monodehydroascorbate reductase (MDHAR), glutathione reductase (GR), peroxidase (POX), superoxide dismutase (SOD), glutathione peroxidase (GPX), catalase (CAT), ascorbate oxidase (AO), and glutathione *S*-transferase (GST), were determined using microplate-based kinetic assays (Aebi, 1984; Murshed et al., 2008).

### Sorghum metabolite profiling

Primary sorghum metabolites were quantified using gas chromatography-mass spectrometry (GC-MS, (Wase et al., 2022). Briefly, metabolites were extracted by methanol: water: chloroform solution from 30±3 mg fresh weight (FW) of frozen samples. Ribitol was added to the extraction solution as internal standard (IS). Dried metabolites were derivatized by methoxyamination and trimethylsylilation and analyzed by a 7200 GC-QTOF system (Agilent, St Clara, CA, USA) using a HP-5MS-UI column (30 m 0.25 mm ID 0.25 μm; Agilent) with 100 x split and splitless modes. Metabolite peaks were annotated by MassHunter Anaysis (Agilent) using the Fiehn Metabolomics Library (Agilent). The 42 most confidently annotated metabolites were quantified by MassHunter Quantitation (Agilent) and the peak heights were background subtracted, normalized by IS and precise fresh weight of the sample to calculate the relative metabolite contents values.

### General characterization of the Arabidopsis dus2 mutants

Arabidopsis Col-0 and *dus2* mutant lines (Line ID: CS857388 and SALK_063052C, https://abrc.osu.edu) were grown under standard Arabidopsis long-day conditions unless otherwise specified (22°C, 60% humidity, and 16h light, 8h dark). All seeds were stratified at 4°C for 48∼72 hours prior to planting. T-DNA insertions were confirmed by PCR using primers designed from Salk T-DNA Express (http://signal.salk.edu/tdnaprimers.2.html; SALK_063052: 5’- GATTGAGCTGGAACGTCTCAC - 3’, 3’- AGCAGAATCTGAGAACCAAATG -5’; CS857388: 5’-ACGCTCTCTTTCAGGATGTTG-3’, 3’-GCTGGTACACCAAGTTTCTCG-5’; L4 primer: 5’-TGATCCATGTAGATTTCCCGGACATGAAG-3’). PCR was performed for 40 cycles with an annealing temperature of 56°C. Germination assays were performed on ½ MS medium with Col-0 and *dus2* mutant seeds at 22°C during each 3-hour interval for 27 hours after seed stratification (Murashige and Skoog, 1962). Two weeks after germination, Col-0 and *dus2* two mutant accessions were divided into separate batches for plant growth rate (kept under long day conditions) and photosynthetic measurements (shifted to a short-day chamber). Four weeks after germination, treatments were applied.

### Measuring impact of abiotic stress on growth rates of Arabidopsis dus2 mutants

Col-0 and two *dus2* mutant lines (CS857388 and SALK_063052C) were used for abiotic stress experiments. Drought treatment occurred at 22°C, with soil water content (SWC) held at 70% for control and 25% for drought set, and lighting period: 8:00 - 21:00 for three days on four-week-old seedlings. Leaf area measurements were performed using RGB imagery after a 3-day water withholding period on plant rosettes using the PlantCV pipeline for trait extraction (Fahlgren et al., 2015). Heat treatment was performed at 37°C for plants grown on soil and media plate. For soil seedlings, 2-week-old seedlings were placed under 22°C and 37°C conditions over 12 days. Plant time-series biomass profiling under these two conditions were performed using Raspi-controlled phenotyping tools (Pheno-Rigs, (Yu et al., 2024a) to collect time-series plant RGB images (imaging-interval: 30 mins, length: 12 days) with quantification of digital biomass of using PlantCV pipelines. For media plate seedlings, 7-day-old young seedling were placed under 22°C and 37°C conditions over 72 hours for observation of seedling color with images.

### Measuring photosynthetic efficiency in Arabidopsis dus2 mutants under abiotic stress

The photosynthetic efficiency of *dus2* and Col-0 lines was evaluated using a gas exchange approach with the LI-6800 (LI-COR Lincoln, NE). Briefly, plants were maintained under short-day (SD) growth conditions (photoperiod: 8:00-17:00, temp: 22°C) to enlarge rosette area to cover the 2cm^2^ small chamber of the LI-6800 device (Brazel et al., 2025). After 4-weeks, plants were kept at 22°C(control), heat (37°C for 24 hours), and drought (25% SWC for 24 hours) for gas exchange measurement. Parameters were set as follows: Flow rate = 400 µmol s⁻¹, Relative humidity of air = 55–60 %, Reference CO_2_ = 400 µmol⁻¹, Mixing fan speed = 10,000 rpm, Heat exchanger temperature: 22°C or 37°C, Chamber pressure = 0.2 kPa, Light source composition 90 % red 10 % blue light, Light intensity = 1000 µmol m⁻^2^ s⁻¹, Measurement time = between 12:00 and 14:00 (Brazel et al., 2025).

### RNA extraction and library construction

RNA extraction was performed using methods from previous studies for both Arabidopsis and sorghum (Burgess, 2023). Approximately five flash-frozen ¼ inch sorghum leaf disks per sample were used as starting material for RNA extractions. RNA extraction was performed using the RNeasy Plant Mini kit (Qiagen #74904) and RNase-free DNase set (Qiagen #79254). Approximately 2.5-10 ug total RNA was used as starting material for mRNA enrichment using the Dynabeads mRNA purification kit (Invitrogen #61006), except with the addition of a second binding and elution step to further enrich mRNA. RNA-seq libraries were then constructed from 10-60 ng mRNA per sample using the Amaryllis Nucleics YourSeq Duet Full Transcript library prep kit with combinatorial dual indexes (now Active Motif #23001 or #23002) according to manufacturer instructions. Prepared libraries were sequenced by Novogene with Illumina NextSeq 500 (PE × 150bp). Arabidopsis RNA was also extracted using Qiagen’s RNeasy kit, per manufacturer’s protocol. Total Arabidopsis RNA was sent to Novogene for mRNA-enrichment and library preparation, then sequenced on Novogene’s NovaSeq X Plus series (PE ×150bp).

### GMUCT sequencing and RNA stability measurement

GMUCT libraries were prepared as previously described (Willmann et al., 2014) using approximately 10 µg of total RNA isolated from the same leaf tissue used for RNA-seq. GMUCT data analyses was performed as previously described (Ganguly et al., 2025), and quality-controlled by ensuring the presence of exon-junction complex footprints in our data. Both RNA-seq reads and GMUCT reads were processed with Hisat2 for mapping and EdgeR to generate RPM-normalize counts (Kim et al., 2019; Chen et al., 2025). mRNA degradation assay was performed as descried (Jia and Le, 2020). Briefly, seven-day-old *Arabidopsis* seedlings of Col-0 and *dus2* were transferred from ½ MS solid medium to ½ MS liquid medium in 12-well culture plates and incubated for 12 hours under 22 °C (control: 8 hours) or 37 °C (heat treatment: 8 hours). Transcriptional inhibitor cordycepin (final concentration = 0.6 mM; Sigma-Aldrich, C3394) was then added to each well for transcriptional arrest. Tissues (∼10 seedlings per biological replicate *3) were collected at 0 h, 15 min, 30 min, 1 h, 2 h, and 6 h after cordycepin treatment for RNA extraction. After the same process of RNA-seq data using Hisat2 alignment and Feature Count pipeline as mentioned above (Liao et al., 2013; Kim et al., 2019), normalization of decay assay was followed as described (Sorenson et al., 2018). Briefly, read counts per gene were normalized based on library size and scaled to 1 million [reads per million (RPM)], further scaled using 30 stable transcripts-derived RNA decay factor to reflect the fact of decreasing total mRNA in the pool during transcriptional arrest (https://rdrr.io/github/reedssorenson/RNAdecay/f/). Decay rate (K) and mRNA half-life were calculated using exponential regression model *y(t)=Ae^−^*^kt^, t_1/2_=ln2/k (A: T_0_ normalized abundance, K: decay rate, t1/2: half-life) and non-linear least square model *y(t)=Ae^−k^t+C*, half-life: t_1/2_= −1/k*ln(0.5-C)/A) (A: T_0_ normalized abundance, K: decay rate, t1/2: half-life, C: residue) based on in house R scripts.

### 5’-6’-Dihydrouridine quantification

Total RNA (1 µg) from *Arabidopsis* leaves was digested to nucleosides using Nuclease PI (0.6 U, Sigma-Aldrich), phosphodiesterase I (0.2 U, Worthington), FastAP (2 U, Thermo Fisher), benzonase (10 U, Sigma-Aldrich), and pentostatin (200 ng, Fisher Scientific) in 25 mM ammonium acetate buffer (pH 7.5). Reactions were incubated overnight at 37 °C and stored at −80 C prior to LC–MS/MS. Digestion enzymes were removed by 75% MeOH precipitation and centrifugation (21,000 g, 10 min); the supernatant was dried and reconstituted in HPLC-grade water. LC–MS/MS was performed on a Waters Acquity UPLC system coupled to a Xevo TQ-XS triple quadrupole mass spectrometer using an HSS-T3 Premier column (1.8 µm, 100 × 2.1 mm). Mobile phases were water + 0.1% formic acid (A) and acetonitrile (B), run with a standard gradient at 0.3 mL min⁻¹. Dihydrouridine was monitored in positive ESI mode (MRM, 247 → 114.9 m/z) with capillary voltage 1 kV, cone 40 V, and desolvation 400 °C. Quantification was based on an external calibration curve using a 5,6-dihydrouridine standard (AdooQ Bioscience).

### Identification and characterization of DHU

Identification of RCMs was performed by intersecting results derived from the High-throughput Annotation of Modified Ribonucleotides (HAMR v1.4) and ModTect (v1.1.5.1) algorithms for sorghum RNA-seq data (Vandivier et al., 2015; Tan et al., 2021). Initially, the paired-end stranded RNA reads were trimmed by Trimmonmatic (v0.32, parameter: TruSeq3-PE.fa:2:30:10:2) to remove adaptors and low-quality reads (Bolger et al., 2014). For HAMR, the forward and reverse reads were mapped to the sorghum reference genome (v3.1.1) using Tophat2 (v2.1.1, --read-mismatches 12 --read-edit-dist 12, --max-hits 10, --lib fr-second strand) (Kim et al., 2013; McCormick et al., 2018), followed by the merging of mapped, removal of multiple-mapped reads, and resolving of splicing alignment using Picard (v3.1.1) and GATK (v3.3.5, (McKenna et al., 2010). Detection were further processed by HAMR with a stringent parameter setting: --min_read_qual 30, --min_read_coverage 50, --hypothesis H_4_, --max_FDR 0.05, which filtered out potential modified sites with a < 50 mapping depth with high-confidence (*FDR* < 0.05) to call modified sites (Vandivier et al., 2015). For ModTect prediction, trimmed reads were mapped to the sorghum genome and associated annotation files using STAR (v 2.7.10b(Dobin and Gingeras, 2015) with the default arguments and ModTect with the following arguments: --minBaseQual 30, --readlength 150. Only the overlapped sites derived from two replicates and two methods were retained as the modified sites for each sorghum sample. The type of RCMs was further classified using a XGBoost-based machine learning model (Palos et al., 2024). The output file of Modtech or HAMR was cross-referenced to the gene annotation files to assign sites to transcripts. Here, the ratio of modified reads over total mapped reads per transcript was calculate as RCM level. Location of DHU sites in sorghum and Arabidopsis dataset were generated following workflow of MetaPlotR (https://github.com/olarerin/metaPlotR). The ±1pb flanking regions of each DHU sites were harvested to calculate the portions of 3nt motifs around DHU sites.

### Population-level genomic analyses in sorghum

Orthologs between Arabidopsis and sorghum were searched based on phytozome12 annotation and confirmed using CoGeBlast (https://genomevolution.org/coge/CoGeBlast.pl, parameters E-value: 1e-5, word_size: 3, matrix: BLOSUM62, TAIR 11 and Sbicolor_454.v3, (Cheng et al., 2017; McCormick et al., 2018). The genomic variants of the six sorghum accessions were identified and annotated using sorghum Association Panel (SAP) data (Boatwright et al., 2022). Briefly, trimmed DNA reads were mapped to the sorghum reference genome (BWA-mem: v0.7.17-r1188, (Li, 2013). Duplicated read-pairs and low-quality mapping in each sample were cleaned using Sambamba (Tarasov et al., 2015) (v1.28.1, mapping_quality >= 1 and not (unmapped or secondary_alignment) and not ([XA] != null or [SA] != null). Variant calling was conducted using the GATK4 HaplotypeCaller function (v4.2.5.00, parameters: --max-alternate-allele:6, --Ploidy:2, (McKenna et al., 2010). The single nucleotide polymorphisms (SNPs) were filtered using GATK hard-filtering (https://gatk.broadinstitute.org). Those high-quality variants were further annotated by variants effect predictor (VEP) using the *S. bicolor* Sbicolor_454.v3 gene annotation (McLaren et al., 2016). For phylogenetic analysis, we retained the bi-allelic SNPs at the four-fold degenerate (4DTV) loci filtered by plink (v1.90b4, --types SNPs -m 2 -M 2) that derived from 365 out of 407 sorghum accessions with complete geographical information (Boatwright et al., 2022). Using the SNP filtering with minor allele frequency (MAF > 0.05, missing data < 0.1) and linkage disequilibrium (LD) prune (plink v1.90b4 parameter: --indep-pairwise 1000 500 0.2, (Purcell et al., 2007), a total of 7,520 SNPs were kept to reconstruct the phylogenetic tree using IQtree (v1.6.12, Parameter: -nt 100 -m MFP, (Nguyen et al., 2015).

### Transcriptome data processing

All RNA-seq reads were trimmed using trimgalore (v0.6.10, -paired, -length 10) and mapped to respective reference genomes using Hisat2 (v2.1.0, default setting, (Krueger, 2015; Kim et al., 2019). FeatureCounts was used for reads quantification (v2.0.1, parameter: -s 1 -p -t mRNA -g ID -O) and further normalized using DESeq2 after the removal of transcripts with less than an average of one count across samples (Liao et al., 2013; Love et al., 2014). For sorghum, time point, accession IDs, and watering conditions were incorporated as the experimental design factors. In Arabidopsis with Condition (Control and Heat) and genotype (Col-0 and *dus2*) included for experimental design Condition-wise differentially abundant transcripts (DATs) over each time point per accession were identified using the cutoff of false discovery rate (*FDR* < 0.05). A log_2_-transformed fold change (FC) > 1 or < - 1was used to filter up (down)-regulated transcripts down-regulated transcripts. Z-score (row-based) normalization was performed to visualize sample-wise heatmap. TPM (transcripts per kilobase million) were used for comparison of transcript abundance across samples for all transcripts.

### Proteomic data processing

#### Proteolytic Digestion

Protein pellets were resuspended in 100 mM Tris-HCl buffer (pH 8.5) containing 4% (w/v) sodium deoxycholate. Proteins were reduced with 10 mM TCEP and alkylated with 40 mM chloroacetamide, then incubated at 45 °C for 5 min with shaking (2000 rpm). Trypsin (in 50 mM ammonium bicarbonate) was added at a 1:100 enzyme-to-protein ratio (w/w), and digestion was carried out overnight at (37 °C with 1500 rpm). After digestion, SDC was removed by phase extraction. Samples were acidified to 1% trifluoroacetic acid and desalted. Peptide concentrations were measured using the Pierce Quantitative Colorimetric Peptide Assay Kit (#23275) following the manufacturer’s instructions.

#### LC-MS/MS for DIA Samples

∼200ng sample was made onto a RSLC column and washed for 5 min with buffer A (www.thermo.com). Bound peptides were then eluted over 35 min onto a RSLC resolving column (gradient: 5%B to 38% B). After the gradient, the column was washed with 90% B for the duration of the run (Buffer A = 99.9% H_2_O, 0.1% formic acid, Buffer B = 80% Acetonitrile, 0.1% formic acid, 19.9% H_2_O, flow rate 400nl/min). Eluted peptides were sprayed into a Q-Exactive HF-X MS (www.thermo.com) for data independent acquisition. Survey scans were taken in the Orbi trap (45000 resolution, determined at m/z 200, mass range: 400-1000m/z. Fixed windows of 20m/z (34 total) with 1m/z overlapping margins were sequentially scanned and fragmented by HCD acquired in the Orbitrap at 15,000 resolutions.

#### DIA Data Analysis

Acquired spectra were processed using DIA-NN (v1.8.1, (Demichev et al., 2020), using the Robust LC (high precision) quantitation strategy with RT-dependent cross-referencing and Deep Learning enabled in library-free mode against Arabidopsis protein sequences available from Uniprot (www.uniprot.org) and search parameters were optimized by DIA-NN, and results filtered at a precursor *FDR* of 1%.

### Nanopore Direct RNA-sequencing (DRS)

Poly(A) tailed RNA was extracted from Arabidopsis wild-type and *dus2* seedlings using 2-3 ug of total cellular RNA or in vitro transcribed transcriptome RNA. Nanopore direct RNA-seq libraries were generated and barcoded following the manufacturer’s instructions (SQK-RNA004). Barcodes were associated with read IDs using SeqTagger (Pryszcz et al., 2025). Raw files were base-called using Dorado (v1.0.0: https://github.com/nanoporetech/dorado) with the following arguments: -v sup, inosine_m6A_2OmeA, 2OmeG,m5C_2OmeC, pseU_2OmeU,--emit-moves, and --min-qscore 10. BAM files were then demultiplexed by filtering for the relevant read IDs from SeqTagger using Samtools and fastq file conversion (Li et al., 2009; Pryszcz et al., 2025). Fastq files were mapped to the *Arabidopsis* transcriptome using Minimap2 (v2.29, parameter: -y, -ax map-ont, and -L (Li, 2018). Nanocompore algorithm was utilized to detect *dus2* dependent modified sites (Leger et al., 2021). First, signal alignment was performed using Uncalled4 (v4.1.0(Kovaka et al., 2021) with signal aligned BAM files used as input to Nanocompore (depleted condition: *dus2*, min coverage: 10, chi-squared q-value < 0.05, GMM log odds ratio > |0.3|).

### Construction of co-expression networks

The co-expression networks were constructed using the WGCNA package with variance stabilizing transformed (VST) normalized read counts with 34,130 transcripts across 96 samples (Langfelder and Horvath, 2008). Initially, a soft threshold (Beta score) was applied to fit the scale-free topology model assumption which generates a minimum number to make the index curve flatten out reaching a high value (>0.85) (Yu et al., 2024b). Further, the adjacency matrix represented by gene expression was transformed into a topological overlap matrix (TOM) to classify genes using a dynamic tree-cut approach. Finally, genes that shared high-confidence correlation (abs(coefficient) > 0.8) were merged into the same module (parameter: deepsplit = 4, tree cut height = 0.2). The pairwise Pearson correlation between each gene from modules and the module-wise eigengene values was tested to characterize each gene’s module membership (MM) under the same module. Lastly, the metabolite profiles, oxidative stress-related enzyme activities, and LI-6800-derived traits were integrated into the network using Pearson correlation between gene expression and these molecular phenotypes. Trait-correlated modules were selected based on their correlation level between MM scores and phenotypes (cutoff: *r* > 0.5, or *r* < -0.5, *p-value* < 0.05). The correlation between gene expression and phenotypes was estimated as gene significance (GS) to interpret the importance of genes related to each molecular phenotype.

### Functional and regulatory network analyses

Functional enrichment analysis of pathways (Kyoto Encyclopedia of Genes and Genomes (KEGG) database) and Gene Ontology (GO) analysis was performed to infer the biological function of genes associated with certain RCMs, transcripts assigned in co-expression modules, and differentially abundant transcripts. ShinyGO was used for GO and KEGG enrichment for both sorghum and Arabidopsis (*FDR* < 0.05, (Ge et al., 2019). The protein-protein interactions (PPI) among genes present in the co-expression network were acquired from the STRING database (parameter: confidence level: > 0.7, (von Mering et al., 2003).

### Data visualization and usage of databases

All statistical visualizations were generated in R (v4.3.2) with multiple packages. Briefly, plots include the line, boxplot, PCA, and enrichment graphs generated by the ggplot2 package, and heatmaps using the pheatmap package (Core Team, 2013). R Scripts used for plots were deposited in (https://rpubs.com/LeonYu/Main-Figures and https://rpubs.com/LeonYu/Supp-Figures). The integrated gene networks PPIs and co-expression derived from WGCNA were generated using Cytoscape (Shannon et al., 2003).

## Results

### Development of a diverse set of sorghum accessions with variation in their molecular and physiological responses to abiotic stress

Phenotypic diversity under drought conditions is ultimately traced to variation in genomic sequence and rewiring of gene expression. Therefore, to capture the genetic and phenotypic diversity present within sorghum, we initially generated a phylogeny from re-sequencing data for 356 out of 400 accessions with demographic information found within the sorghum association panel (SAP, (Boatwright et al., 2022)**Figure 1A**). Cross-referencing this phylogeny with the SAP field trial (in Arizona in 2019) under standard conditions, we selected six accessions to comprise a small but diverse panel based on seed viability, genetic distinctiveness, and phenotypic variation (shoot and root biomass, plant height, and yield). Among the selected lines, four accessions (A, B, E, and F) are African accessions, and the other two accessions (C and D) are U.S. breeding accessions (**Figure 1A**). Variant calling and annotation of variants were performed to examine the genomic variation between these accessions. In total, 8,725,449 variants were identified among the six accessions relative to the published reference genome, with more than 50% of variants located in intergenic regions, followed by upstream and downstream gene regulatory regions (21.0% and 19.5%, respectively), intronic regions (5.61%), and splicing regions (0.23%; **Figure S1, left panel**). Approximately 10-20% of the genomic variation in the panel was comprised of unique SNPs and INDELs specific to each genotype (**Figure S1 right panel**). In summary, these data suggest that we were successful in selecting six distinct accessions, representative of the genetic diversity within the species, and likely the wide range of environments in which sorghum can grow.

**Figure 1.**
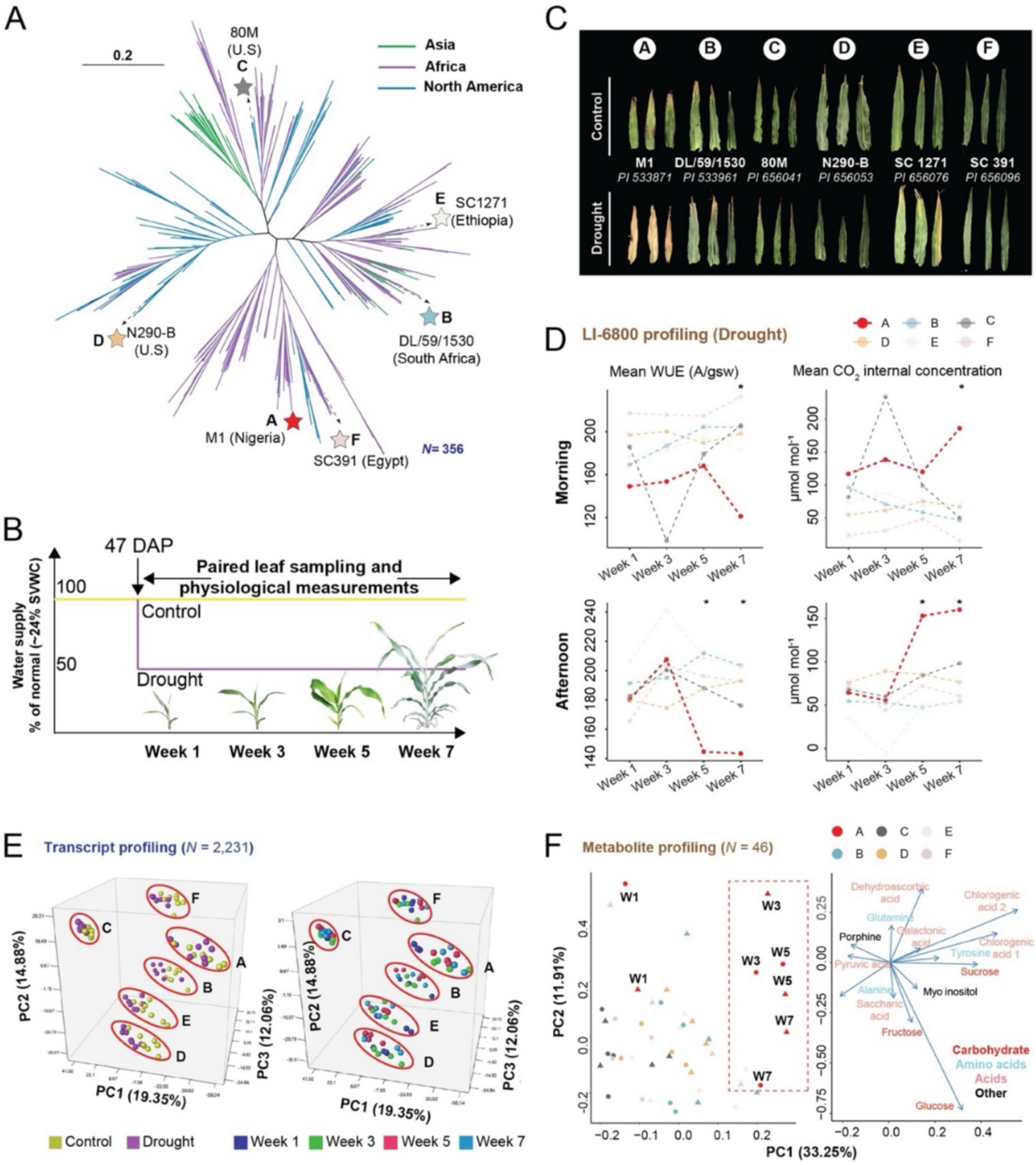
Drought tolerance diversity of selected sorghum genotypes. **(A)** Reconstruction of an unrooted phylogeny of the Sorghum Association Panel (SAP). The six sorghum accessions used in this study are marked along with the geographical locations colored to represent African (green), Asian(purple), and North American (blue) regions. The placement of the six accessions used in this study is denoted with different colored stars that match the color scheme throughout this work. (**B)** Six sorghum accessions were grown at the Arizona in the field with treatment (reduced water supply) imposed 47 days after planting (DAP). Paired molecular sampling (RNA-seq, metabolites, and oxidative stress chemicals), were collected alongside physiological measurements (e.g., LI-6800 photosynthesis-related traits). **(C)** Representative leaf tissue collected at the terminal stage of sampling in the field is shown. **(D)** Line graphs denote the mean WUEi and internal CO_2_ concentration values measured during the morning (top) and afternoon (bottom) at each time point for drought-treated plants (*N* = 5 per accession). Asterisks indicate significant differences of parameters derived from the comparison of each accession relative to “A” (*p* < 0.05). **(E)** Two principal component analyses (PCAs), based on the top 10% most abundant transcripts, highlight the separation caused by genotypic and treatment effects (left) and genotypic and temporal effects (right). **(F)** The left PCA displays separation based on treatment (triangle = drought; circle = control) and genotype (colors similar to those in panel C). The right PCA loading plot highlights major contributors to the left PCA, with the length of the line denoting the contribution of a particular molecule to the PCA.

To assess the performance of this panel under heat and drought conditions, we grew the six accessions in replicated plots as part of a large field trial conducted during the summer of 2020. All plants were well-watered (WW) until beginning of the reproductive stage (appearance of the flag leaf; 47 days after planting; **Figure 1B**), after which irrigation was reduced for a subset of plots to mimic drought treatment. Phenotypic measurements and molecular sampling (for transcriptomic, metabolomic, and biochemical traits) occurred during the 1_st_, 3_rd_, 5_th_, and 7_th_ week after the beginning of the drought treatment (**Figure 1B**), with destructive measurements (e.g., biomass and root microbiome composition) occurring at the end of the season. Treatment effects were readily apparent at the end of the season (**Figures 1C**), with the M1 (“A”) accession showing the most pronounced leaf necrosis. Leaf level physiological traits, such as water use efficiency (WUE) and CO_2_ net assimilation (A_net_) exhibited similar results, with the “A” accession displaying hallmarks of water stress, particularly at the later time points (week 5 and week 7, **Figure 1D**), whereas other accessions, such as “B” and “C”, showed a more robust response. Thus, in addition to genotypic diversity, these data suggest that these six accessions displayed phenotypic diversity to drought stress and may provide information on the molecular mechanisms by which sorghum responds to water limiting conditions.

### Sorghum accessions display variable molecular responses to stress

To characterize the molecular responses to water limiting conditions in our panel, we next assessed the change in transcript abundance over the two months of the treatment (Weeks 1, 3, 5, and 7) using RNA-seq. Initial quality control revealed high correlation between replicates (Pearson Correlation; r > 0.95, **Table S1**). To unpack the transcriptomic responses to treatment in the six accessions, we examined the top 10% of most variable transcripts based on the standard deviation (SD) of the global transcript abundance for principal component analysis (PCA). This approach uncovered strong genotypic and treatment effects represented by both PC1 and PC2, with accession “C” exhibiting the least separation due to stress or time, and accession “A” exhibiting the most (PC1 = 19.35%, PC2 = 14.88%, **Figure 1E**).

To gather additional clues as to how these accessions responded to drought treatment at the molecular level, we next examined the stress-induced changes in metabolites and oxidative stress markers in sorghum leaves (Das and Roychoudhury, 2014; Obata et al., 2015). Metabolites and oxidative stress markers were measured on leaf samples collected simultaneously with those used for RNA-seq analyses, with GC-MS-based quantification and annotation of primary metabolites and spectrophotometric-based quantification of oxidative stress markers. A PCA of global changes in the leaf primary metabolome (*N* = 46) across our panel revealed that, as for the transcriptome, genotype explained much of the variation between samples (**Figure 1F**, left panel; PC1: 33.25 %). Interestingly, samples associated with accession “A”, weeks 3, 5, and 7 (both conditions), separated from all other samples, and this separation was largely driven by levels of the three main soluble sugars (sucrose, glucose and fructose) and chlorogenic acid (**Figure 1F**, right panel). Overall, we identified strong genotypic effects for both visible and molecular phenotypes. As expected for this species, most accessions displayed resilience under stressed conditions, whereas accession “A” performed poorly in under stress conditions. Thus, this panel presents a suitable comparative framework to examine transcriptional and post-transcriptional regulators of drought and heat stress responses in sorghum under field conditions.

### Integrated network analysis uncovers a link between DHU and photosynthesis traits

With each of these molecular and physiological traits examined, we next took a comparative systems-level approach to connect these traits to their molecular pathways and putative regulators. To do so, we constructed a co-expression network starting with the top 10% of most variable transcripts, as measured by median absolute deviation (MAD) score. We then recursively increased the number of included transcripts until we achieved an optimized network with the fewest number of unassigned transcripts and the highest scale-free topology model (**Table S2**). The resulting network contained 22,314 transcripts classified into 29 modules with a low proportion of unclassified transcripts (*N =* 2,982) and high scale-free model correlation (*r* = 0.91, slope = -1.93, **Table S2**). Using this framework, we incorporated phenotypic data into modules using trait-module correlation. Briefly, the first principal component value (eigenvalue) per module was calculated using their transcript abundance PCA. Leaf-level physiological measurements (LI-6800), metabolite abundances, and oxidative stress marker abundances were then correlated with the eigenvalues of each module (**Table S3**), leading to 19 modules with significant trait-module correlations (*p* < 0.05, **Figure 2A**, left panel). Several modules (e.g. “blue”, “greenyellow”, “pink”, and “purple”) were strongly correlated with multiple traits. For instance, “blue”, “purple”, and “pink” modules are closely related to most photosynthesis traits (> 90% LI-6800 entries), whereas “purple” and “greenyellow” modules are highly correlated with many metabolic traits and oxidative stress chemical abundances (**Figure 2A**).

**Figure 2.**
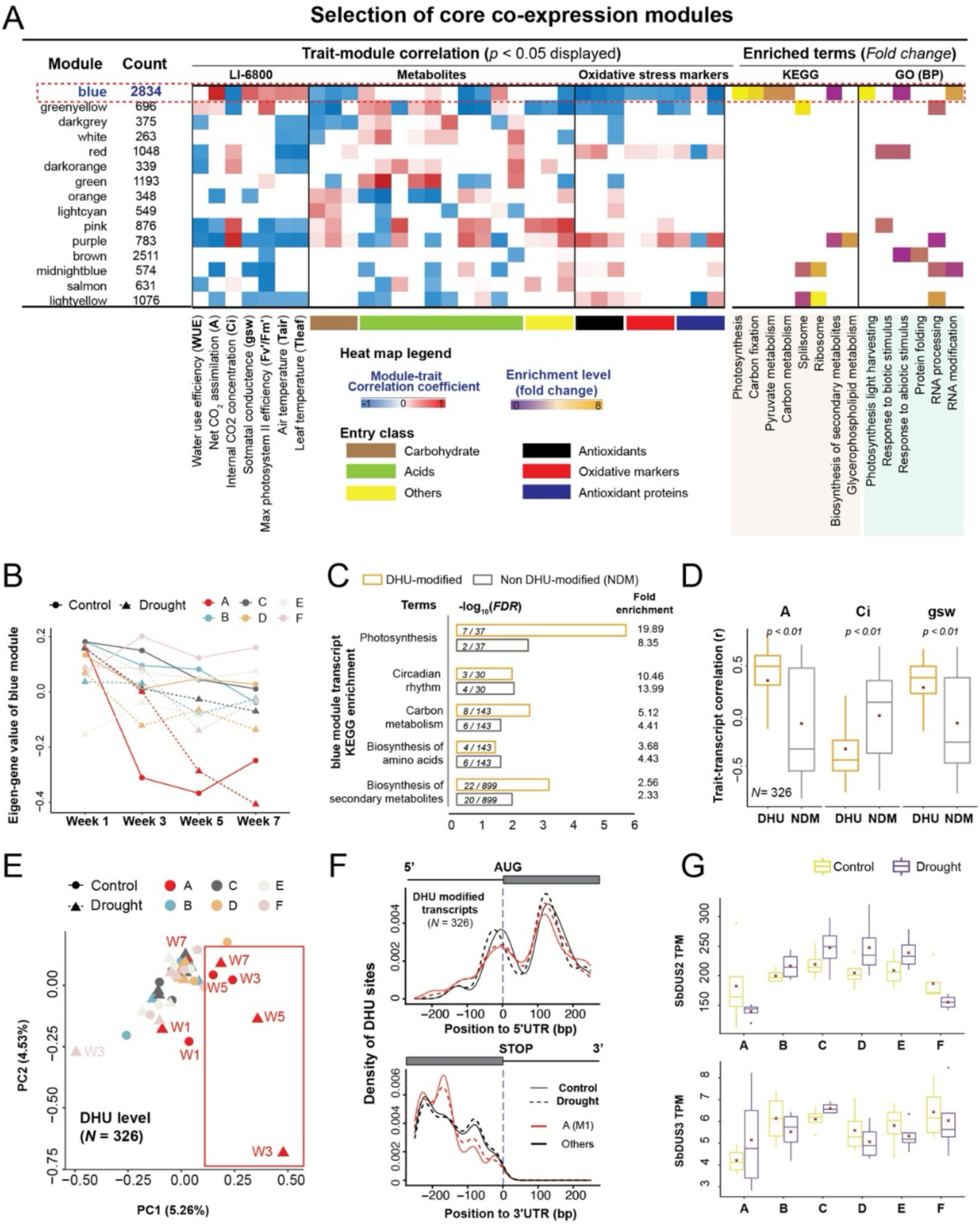
Selection of co-expression modules contributing to sorghum drought tolerance. **(A)** The number of transcripts per module, correlation between traits and the module (blue and red scale), levels of KEGG and GO enrichment levels (purple and brown scale), and the known proteins related to RCM recognition, deposition, and removal located in each co-expression module. Scales for module-trait correlation values (blue to red scale: -1 to 1), as well as GO-enrichment levels (purple to yellow scale: fold changes) are shown to the bottom left. Only those module-trait correlations and enrichment levels with a *p* < 0.05 are shown. **(B)** The eigenvalue of the photosynthetic module, divided by accession, treatment, and time, was plotted. (**C)** Correlation of transcript abundance and modification was compared between 326 transcripts been modified by both DHU and other RCMs in the blue module. **(D)** Bar plots denote the log_2_(*FDR*) score of consensus enrichment terms, with number of genes associated with the term over the total genes in the term (middle), and the fold change for each term (right) **(E):** Box plots denote the correlation coefficient (r) between transcript abundance and each photosynthetic trait (A, Ci, and gsw) between DHU-modified group and the non-modified group. **(E)** Principal component analysis (PCA) of DHU-modified transcripts from the “blue” module (*N =* 326 transcripts). **(F)** Metaplot shows density of DHU sites across upstream (downstream) 250bp genomic region near start (stop) codon over 325 transcripts in accession “A” and other five accessions under control and drought conditions. (**G)** TPM of *SbDUS2* and *SbDUS3* across the two conditions and six accessions.

To gain insights into the biological processes associated with modules, module-wise GO and KEGG enrichment analyses were performed (modules with *FDR* < 0.05 are displayed, **Figure 2A**, top panel). This approach revealed modules with biological processes of interest, such as stress response (“blue”, “red”, and “pink”), RNA processing and modification pathways (“blue”, “greenyellow”, “purple”, “midnightblue”, and “lightyellow”), and photosynthesis terms (“blue”, **Figure 2A and Table S4**). Specifically, in addition to the observed module-trait correlation, the “blue” module was strongly enriched for transcripts associated with photosynthesis traits (eg: net CO_2_ assimilation: r = 0.72) based on both GO and KEGG terms (fold change > 5, *FDR* < 0.05, **Table S4**). Interestingly, an examination of eigengene values within this module revealed a strong genotypic effect, where transcripts derived from the “A” accession cluster separately from the other accessions (**Figure 2B**), and therefore may explain the physiological distinctiveness of this accession. Given the strong trait module correlations and the over-representation of stress and photosynthesis associated genes, the “blue” module thus represents a strong link between phenotypic and molecular responses to stress treatment.

A close examination of genes with strong module membership (kME; **Table S5**) in the “blue” module revealed two genes encoding putative dihydrouridine synthases (DUS), which convert uridine to dihydrouridine (DHU; Sobic.006G158100 and Sobic.006G038500). A phylogenetic analysis using a maximum likelihood tree derived from an amino acid sequence alignment of human, fission yeast, budding yeast, and multiple plant DUS proteins suggests that Sobic.006G158100 is more similar to eukaryotic DUS2 (**Figure S2**; SbDUS2), whereas Sobic.006G038500 fell into the DUS3 clade (**Figure S2**; SbDUS3). A third sorghum DUS, Sobic.010G262000, fell into the DUS1 clade but was not found to be associated with plant performance traits within the “blue” module (SbDUS1). The strong association between these two genes and the blue module implies that their transcriptional regulation, and potentially their function, is associated with plant performance under stress.

Given the inclusion of DHU synthase in the stress and photosynthesis-related “blue” module, we examined the RNA-seq data for hallmarks of RCMs that impact the Watson-Crick-Franklin base-pairing edge. DHU belongs to a subset of RCMs that interfere with RNA-seq library preparation (specifically the reverse transcription step), leaving patterns of mis-incorporated nucleotides that can be assigned to specific modification classes bioinformatically (Vandivier et al., 2015; Helm and Motorin, 2017; Tan et al., 2021). We thus processed our RNA-seq data through two independent RCM predictors (HAMR and Modtect) and took the intersection of called sites (**Table S6**), resulting in 20,096 high-confidence RCMs associated with 3,715 transcripts among the six main classes of HAMR/Modtect identifiable RCMs (**Table S7**). Of these, a total of 3,008 DHU sites on 1,370 transcripts were detected, with 326 transcripts associated with predicted DHU modification residing within the “blue” module (**Table S8**). To determine if DHU is preferentially associated with photosynthesis and stress related transcripts in this module, we performed KEGG enrichment of these 326 transcripts and a similar number of non DHU-modified (NDM) transcripts within the module as background (random selection bootstrap = 10). This approach uncovered an elevated enrichment level of “photosynthesis” terms in the DHU-modified sets compared with the “non DHU modified” set (*FDR* < 0.05, **Figure 2C**). Lastly, to determine if there was a relationship between DHU abundance and field-derived traits, a pair-wise Pearson correlation was performed between transcript abundance and photosynthetic trait measurements (A, Ci, and gsw, respectively) for all DHU-modified and a similar number of randomly selected non-DHU modified transcripts (**Figure 2D**). This analysis revealed a strong correlation between transcripts associated with DHU modification and photosynthetic traits, suggesting a close relationship between DHU modification status, transcript abundance, and photosynthetic processes in sorghum.

We next assessed whether the transcriptomic accession-associated variation was also observed in the DHU deposition on these 326 modified transcripts. A PCA revealed that the DHU status for these transcripts in the “A” accession was markedly different at most time points and conditions relative to the other accessions (PC1 was primarily explained by DHU variation in the “A” accession; **Figure 2E**). We explored this accession-driven variation further by determining if the pattern of DHU deposition on these transcripts was different in the “A” accession relative to the others. Based on a metaplot profiling of DHU deposition on the 326 DHU-modified transcripts, we observed variation at both the 5’ and 3’ ends, with the most dramatic variation near the 3’ end of the CDS in the “A” accession relative to the others (**Figure 3F**). In sum, these data highlight the association between DHU status and photosynthesis, suggesting that variation in this molecular phenomenon between accessions be linked to plant performance.

**Figure 3.**
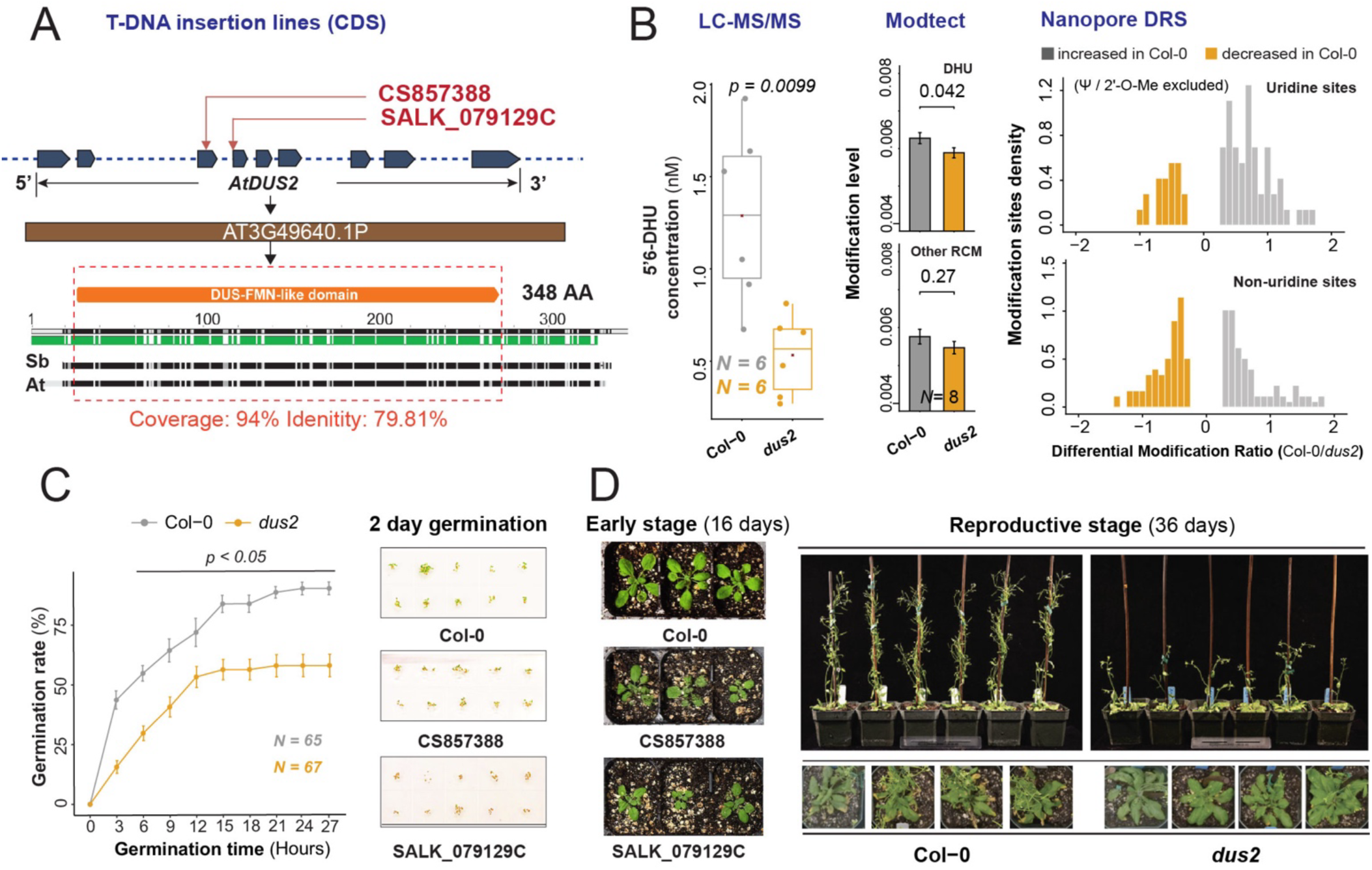
Characterization of the *dus2* mutant in Arabidopsis. **(A)** Schematic of the *AtDUS2* locus (top) with the T-DNA insertion sites in the third and fourth exon denoted. A protein alignment of the Arabidopsis and sorghum DUS2 proteins is depicted below, with the DUS-FMN-like domain outlined in orange. Alignment coverage and amino acid identity is shown below. **(B)** Left: Comparison of 5,6-dihydrouridine between two genetic backgrounds using LC–MS/MS analysis of hydrolyzed total RNA with external calibration using synthetic 5,6-dihydrouridine standards under (10, 50, 100 nM); Middle: Density plot of significantly modified sites (q-value < 0.05) between Col-0 and *dus2* of uridine-associated/non-uridine associated sites using Nanopore DRS**. (C)** Quantification based on imaging of seeds grown on MS plates, with each datapoint representing three plates. Germination was calculated as the full emergence of cotyledons with images after 2-day germination**(D)**Top-view images of two independent dus2 T-DNA insertion lines at 2-week of growth (left). Side images of the two backgrounds at the point of silique maturation for Col-0 (36 days; top), and top-view images of the rosettes captured at the same timepoint (bottom).

### SbDUS2 is associated with DHU deposition on photosynthesis-related transcripts in sorghum under abiotic stress

Because both SbDUS2 and SbDUS3 appeared in this DHU-associated module, we examined the reproducibility of transcriptomic interactions within the “blue” module to help guide the selection of a candidate for subsequent studies. To do so, we first developed a child network of all transcripts in the “blue” module that shared high confidence edges with *SbDUS2* and *SbDUS3* and were DHU modified (*N* = 29**, Figure S3A and Table S5**) and examined the relationship of these transcripts with both *SbDUS2* and *SbDUS3* transcripts and DHU abundance. Interestingly, we observed a negative correlation between DHU status and the abundance of these 29 interacting transcripts in our field data (**Figure S3B**). We then examined the abundance of each of these transcripts in a publicly available sorghum tissue atlas dataset (accession: BT×623; (McCormick et al., 2018). Not only where these 29 DHU-modified transcripts most highly abundant at similar developmental stages in the sorghum tissue atlas as in our dataset, we also observed a strong negative correlation between DHU levels and 11 out of the 29 focal transcripts (*p* < 0.05; **Figure S3C**). Notably, *SbDUS2* and *SbDUS3* were also predominantly expressed in leaf tissues, and largely during late vegetative and flowering developmental stages, similar to our observations in our six accessions (**Figure S3D**). We then asked which SbDUS most likely influences DHU deposition on mRNAs within the “blue” module by assessing the correlation between the score of the first principal component (PC1) of 326 transcripts-derived DHU PCA **(Figure 2E)** and the transcript abundance of *SbDUS2* and *SbDUS3* (**Figure S3E).** This analysis revealed a stronger correlation between *SbDUS2* and PC1 scores (*r* = -0.63, *p* = 1.4e-06) than for *SbDUS3 (r* = -0.39, *p* = 0.0064), implying that SbDUS2 might be more tightly associated with DHU status. Additionally, *SbDUS2* exhibited significantly higher transcript abundance (TPM) in both datasets (*p* < 0.0001, **Figure S3D and Figure 2G**). Not only was *SbDUS2* on average ∼40x more abundant than *SbDUS3* across the six accessions and in the sorghum tissue atlas data, Sb*DUS2* also displayed an overall decreased transcript abundance under water limiting conditions in the “A” and “F” accessions and an increase in all other accessions (**Figure 2G**), whereas *SbDUS3* displayed no significant treatment effect. We also assessed *SbDUS1*, which was more highly abundant than *SbDUS3*, but did not associate with transcripts in the “blue” module or performance-related traits and displayed minimal transcript-level differences in response to treatment (**Figure S3F**). Thus, based on its consistently high transcript abundance across two datasets and its stress-response pattern, we focused analysis on DUS2 and its potential contribution to DHU modification under abiotic stress conditions.

### DUS2 mutants display DHU level reduction associated with growth and developmental delays in Arabidopsis

Convinced that DHU was somehow linked to photosynthetic traits in sorghum, and that SbDUS2 was the likely enzyme responsible for DHU deposition on photosynthesis-associated transcripts, we next turned to the genetic model, *Arabidopsis thaliana*, to gain a better understanding of how DHU might impact plant performance. Based on our phylogenetic assessment of the DUS1/2/3 protein family across eukaryotes, we identified the Arabidopsis *Dus2* ortholog, which encodes a protein that is 79.81% identical to SbDUS2 at the amino acid level (**Figures S2 and 3A**). We obtained two T-DNA insertion lines that disrupted the third and fourth exons, respectively, within the coding region of *AtDus2* (AT3G49640, **Figure 3A**, CS857388 and SALK_079129C, https://abrc.osu.edu/). We confirmed that these insertions disrupted *AtDUS2* transcription using both Illumina and Oxford direct RNA-seq (ONT-DRS; **Figure S4A**). Loss of *AtDus2* resulted in a significant reduction of DHU based on LC-MS/MS of total RNA extracted from mature rosette leaves (**Figure 3B; left, Table S9**), as well as a significantly lower global HAMR/Modtect-derived DHU level not observed for other RCMs detectable by these algorithms (**Figure 3B; middle,** *N* = 8). Although a DHU-aware base-calling algorithm is not currently available for ONT-DRS, as another layer of cross-validation, we used ONT-DRS to detect differentially “modified” U sites between Col-0 and the *dus2* background that were not associated with the two currently detectable Uracil modifications, Ψ or 2’-O-Me-U (**Figure 3B; right**). After excluding these modifications, we observed an increase in modified U sites in Col-0 relative to the *dus2* background, suggesting that these modified sites are AtDUS2 dependent. Based on HAMR/Modtect-derived data in both Arabidopsis (Col-0) and sorghum samples, we observed DHU being deposited in a similar U-tract sequence context within mRNAs as was observed in yeast (**Figure S5A**), with a preference for the CDS over UTRs (**Figure S5B**, Finet et al., 2022b). These data suggest that DHU is found within Arabidopsis mRNAs, and that at least a subset of them are dependent on *AtDUS2*.

In addition to diminished DHU levels, Arabidopsis *dus2* mutants displayed reduced germination (**Figure 3C**; *p <* 0.05, *N* > 50) and slower growth rates, as well as shorter stature and increased time to flower in both *Atdus2* mutant lines relative to wild-type controls (Col-0, **Figure 3D and Figure S4C and S4E**), although the *Atdus2* mutants eventually reach normal height relative to Col-0 (**Figure S4F**). Ultimately, these phenotypes led to an extended period of vegetative growth and delayed leaf senescence in *Atdus2* relative to Col-0 (**Figure 3D**), confirming that loss of AtDUS2 leads to abnormal growth and development under control conditions.

Given the importance of DHU for function of other classes of RNAs, we next assessed whether the *Atdus2* mutant background exhibited an mRNA and protein-level response that might explain the observed delayed growth phenotype. To explore transcriptomic and proteomic responses, we compared profiles from mature rosette leaves of *Atdus2* mutants versus Col-0 controls. Illumina RNA-seq was used to determine which transcripts were differentially abundant and untargeted LC-MS/MS was used to determine if protein abundance was impaired by loss of *AtDus2*. Loss of *AtDus2* led to 807 and 1,096 transcripts with increased and decreased abundance, respectively (*FDR* < 0.05, |log_2_FC| > 1; **Figure S6A, Table S10**). At the protein level, 2,901 proteins with reproducible detection across replicates (*N* = 3 per genotype) were examined for changes in abundance between *Atdus2* and Col-0. We found 128 and 147 proteins with increased or decreased abundance (*padj* < 0.05; **Figure S6B**), respectively, in the mutant, with a strong positive correlation in the change between genotypes between transcript and protein abundance observed for those where matching data were available (*N* = 275, *r* = 0.43 and *p* = 1.7e-13, **Figure S6C**). We then integrated the status of RNA, DHU, and protein abundance in the two genetic backgrounds, focusing on a set of 38 DHU modified transcripts that displayed differential abundance at all three levels between Col-0 and *Atdus2* backgrounds (**Table S11**). This subset displayed a strong positive correlation between transcript and protein abundance (*r* = 0.54, *p* = 0.00047; **Figure S6D**), and a negative correlation between DHU levels and protein abundance (*r* = -0.35, *p* = 0.038; **Figure S6E**). These data suggest that DHU is predominantly, but not always, negatively associated with mRNA and protein abundance.

### AtDus2 loss-of-function leads to alterations in mRNA stability

Given the observed negative relationship between DHU level and mRNA abundance in both Arabidopsis and Sorghum, we next assessed whether DHU was impacting RNA stability. We first examined differences in global uncapped mRNA profiles between Col-0 and *Atdus2* using the genome-wide uncapped and cleaved transcripts (GMUCT) approach (Gregory et al., 2008; Willmann et al., 2014). In this framework, a higher proportion of uncapped transcripts (GMUCT reads/ RNA-seq reads) indicates reduced mRNA stability, whereas a lower proportion suggests enhanced RNA stability. To assess data quality, we ensured the captured of exon-junction complex footprints in our libraries for both genotypes (Lee et al., 2020). Using HAMR/Modtect detection, we identified a total of 426 DHU-modified transcripts in Col-0 under control condition, and 302 of these transcripts were associated with reduced DHU levels in the *Atdus2* background (**Table S12**), suggesting their DHU status was *AtDUS2* dependent. These DHU-level dependent transcripts were not only more abundant at transcript level in the *Atdus2* background relative to non DHU-modified transcripts based on cumulative frequency (**Figure S7A,** *p* < 1e-16, solid lines), GMUCT also uncovered a significant reduction in the proportion of uncapped RNAs in the *Atdus2* background relative to Col-0 (**Figure 4A**, blue boxplots, *p* = 0.00087). This reduction was not observed for non DHU-modified transcripts (**Figure 4A**, red boxplots, *p* = 0.25), suggesting that the presence of DHU on these transcripts in a Col-0 background was influencing their degradation.

**Figure 4.**
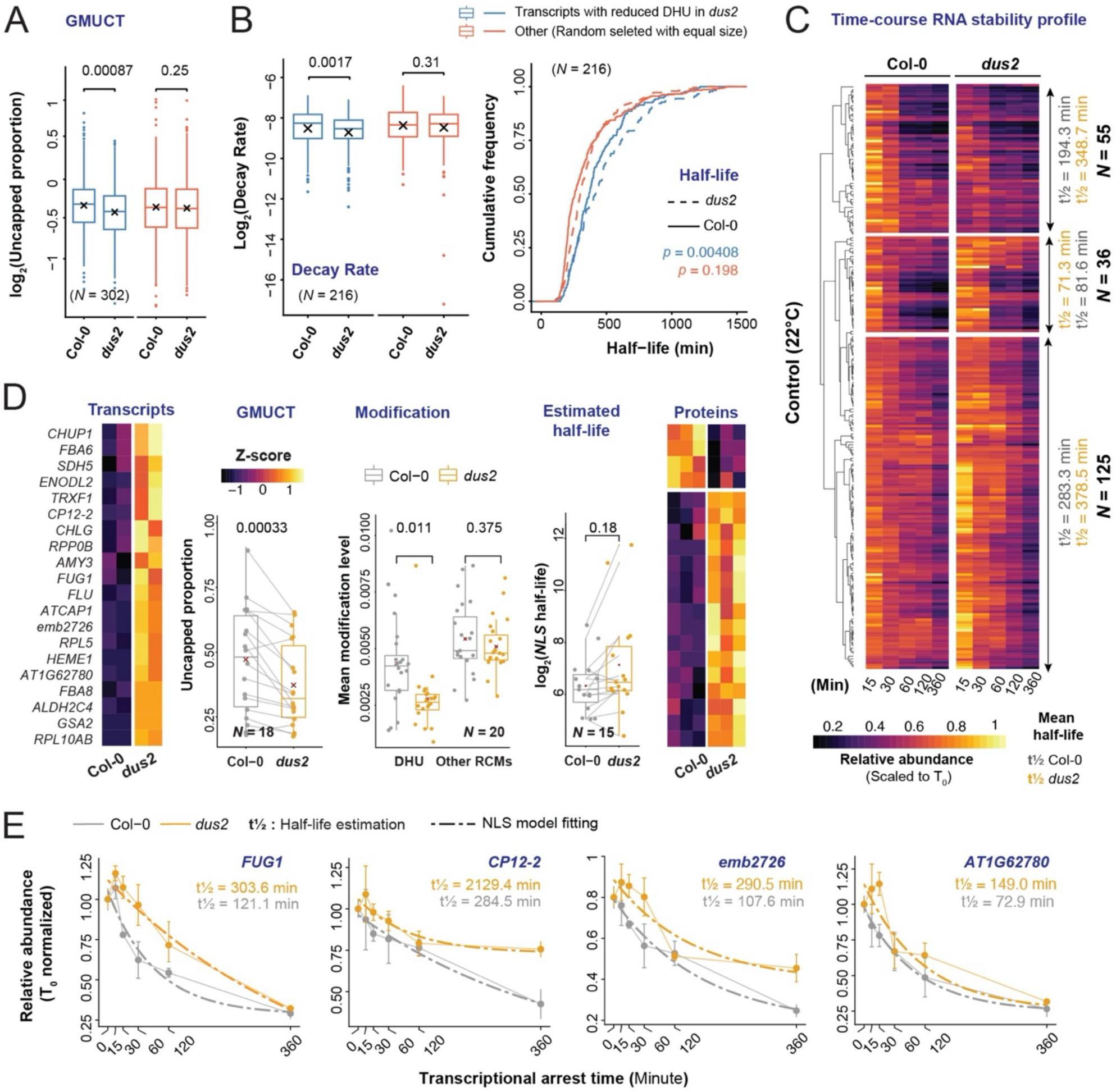
Altered RNA stability between Col-0 and *dus2* background. **(A)** Boxplot shows comparison of GMUCT-derived uncapped portions per transcripts over transcripts associated with lower DHU level in *dus2* (blue boxplots) and other transcripts (red boxplots) in Col-0 and *dus2* background, examinate by Wilcox test. **(B)** Decay rate (*K*) on groups of transcripts with lower DHU level in *dus2* mutants (blue boxplots) were compared with subset of randomly selected transcripts with equal sample size (red boxplots) using t-test (*p* < 0.05) on the left; Cumulative frequency of estimated half-life of transcripts in Col-0 (solid line) and *dus2* (dashed line) background with lower DHU level in *dus2* mutants (blue lines) were compared with randomly selected transcripts (red lines) on the right. **(C)** Heatmaps show differentiated decay rate of transcripts derived from panel (B), which majority faster decay in Col-0, partial similar decay between two genotypes, and small groups with slower decay in Col-0 under both control and heat conditions. **(D)** Heatmap of 20 transcripts associated with differential DHU, transits abundance, and protein abundance was plotted. Boxplots show GMUCT and RNA modification level of 16 transcripts with reduced protein in Col-0 relative to *dus2* (Wilcox test, *p* < 0.05), along with estimated half-life comparison of14/16 transcripts (Wilcox test). **(E)** RNA decay curve 6-hour experiment and line graph (error bars displayed) of four selected transcripts associated with higher DHU level and faster degradation in Col-0 than *dus2*, with estimated half-life were using NLS model.

To further test this model, we performed a time course transcriptional arrest experiment, treating 7 day-old Arabidopsis seedlings with the transcription inhibitor, cordycepin over a period of six hours (**Figure S7B**). After QC (correlation among replicates > 0.9, aligmnet rate > 90%) and data normalization based on (Sorenson et al., 2018), including the removal of low-abundance transcripts at T_0_ and transcripts with high standard error (T_0_ average TPM > 8 and SE < 0.5), we retained 12,450 trancripts for modeling using both an exponential regression model (*Exp*) and a nonlinear least squared model (NLS) to estimate the decay rate (*K*) and half-life (min) for each transcript (**Supplementary Table 13**). Using either model, we did not observe a significant change in global RNA decay rates (**Supplementary Table 13**). However, when examining just those transcripts for which we could assign DHU status and model decay rates (216/302 transcripts with reduced DHU in *dus2*), we saw both a significant decrease in decay rates and corresponding increase in mRNA halflives (**Figure 4B**; left = decay rate, right = half life estimation). These 216 DHU-modified transcripts could be further divided into three groups based on their half-life patterns using clustering as a reference (**Figure 4C and Figure S8A,** average half-lives denoted for each cluster), with a great portion all displaying a longer half-life in the *Atdus2* background (*N* = 125; **Figure 4C**). Interestingly, these more stable transcripts were predominantly associated with photosynthetic and translation-related GO-terms (**Figure S8B, top**). These data suggest that, in agreement with the GMUCT data, DHU is predominantly marking unstable or degrading transcripts.

We next returned to the 38 transcripts for which we saw links between DHU, transcript, and protein abundance (**Figure S6D and S6E**). Of these, 20 showed both increased transcription and significantly reduced DHU levels in the *Atdus2* background (*p* = 0.011; **Figure 4D middle**) with 16 out of these 20 displaying significantly increased protein abundance. As expected, these transcripts displayed limited variation in non-DHU RCMs detected by HAMR/Modtect (*p* = 0.375). These differentially DHU-modified transcripts are involved in KEGG-defined metabolic pathways, ribosome function, and photosynthesis-related processes, with most (14/20) localized to either the chloroplast or mitochondria, although these transcripts originate from the nuclear genome (**Table S11**). A majority of these DHU modified transcripts with sufficient GMUCT coverage (15/18) also displayed a significant decrease in the proportion of uncapped reads relative to Col-0 (*p*= 0.00033, **Figure 4D middle**), suggesting DHU-dependent transcript degradation. Half-life modeling largely agreed with this model of DHU-dependent degradation, as 12/15 transcripts (with sufficient data to model) showed an increased half-life in the *Atdus2* background, with half-life estimates increasing by ∼2-10 fold for four transcripts as displayed (**Figures 4E**, full dataset and modeled rates found in **Supplementary Table 13**). Thus, under normal growth conditions, DUS2, and the RCM DHU, appear to be important for turnover of a core subset of transcripts associated with photosynthesis and metabolism.

### DHU impacts photosynthesis-associated traits under abiotic stress in Arabidopsis

Given SbDUS2’s connection to performance under abiotic stress in sorghum, we next sought to validate this role in the context of a *Atdus2* loss-of-function background in Arabidopsis plants. First, we assessed whether there was a DUS2-dependent impact on plant performance under the two main stressors sorghum experienced in the field, water deficit and heat stress, applied separately here for Arabidopsis. We monitored vegetative growth and development of Col-0 and *Atdus2* mutants seedling (SALK_079129C and CS857388) under water withholding or heat stress conditions. Water withholding was performed by reducing soil water content (SWC) from 70% (control conditions) down to 25% (drought) over six days and then holding SWC at 25% for three additional days on four-week-old seedlings, with images been analyzed to estimate approximate growth rates (**Figures 5A and S4B**). As expected, Col-0 plants displayed reduced rosette area under water withholding conditions, indicating that the stress was successfully applied to these plants (**Figure 5B**). Plants lacking *Atdus2* showed a significant ∼50% reduction in rosette area relative to Col-0 under both control and water withholding conditions (**Figures 5B** and **S4C**). Heat stress, applied to two-week old seedlings for 12 days, resulted in a similar relative decrease in plant growth for both Col-0 and *Atdus2* mutants, however, the *Atdus2* mutant backgrounds showed significantly reduced survival rate at the end of the experiment (**Figures 5C** and **5D**). Seven-day-old *Atdus2* mutant seedlings showed a similar response when exposed to heat stress for 72 hours (37°C) on ½ MS plates, suggesting this is a robust sensitivity to heat stress. We next assessed photosynthetic capacity via the LI-6800 system (net CO_2_ assimilation) in both the Col-0 and *Atdus2* drought and heat stressed plants (**Figure 5E**). Interestingly, we observed a slight, but non-significant decrease in net CO2 assimilation under control conditions between *Atdus2* and Col-0, whereas the *Atdus2* background showed a significant reduction under both stress conditions (*p* <<< 0.05 in both treatments; **Figure 5E**). These data suggest that AtDUS2, and DHU, is important for full photosynthetic capacity under drought and heat stress.

**Figure 5.**
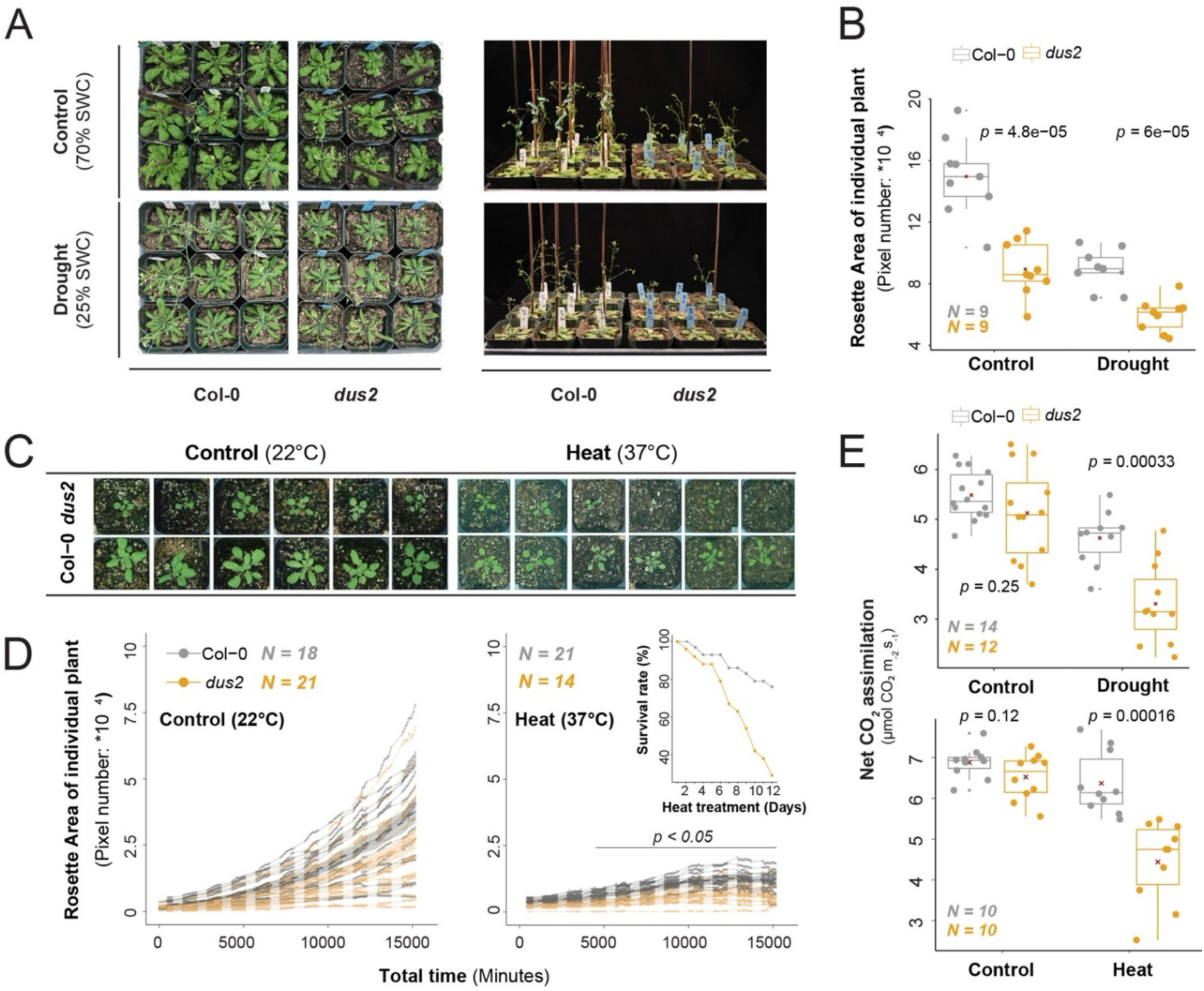
Examination of abiotic stress response between Col-0 and *dus2* in Arabidopsis. **(A)** Arabidopsis rosettes and seedling architecture of 4-week-old seedlings among control (70% SWC) and drought (25% SWC) conditions were displayed. **(B)** Comparison of Arabidopsis rosette area at the last day of drought treatment between Col-0 and *dus2* plants under 70% SWC and 25% SWC conditions (t-test, *p* < 0.05). **(C)** Images show Arabidopsis rosette characteristics at the 12^th^ day of heat treatment between two genotypes. **(D)** Image-based phenotyping of Arabidopsis rosette area over 12 days of vegetative growth under control and heat conditions (30 minutes interval) processed by PlantCV (Yu et al., 2024a). Inset panel under heat depicts survival rates over time.**(E)**Photosynthetic efficiency (net CO_2_ assimilation) was measured on the largest rosette leaf using a LI-6800 system. CO_2_ net assimilation rate under heat (24h at 37°C) and control (22°C), control (70% SWC) and drought (25% SWC) were measured with more than 10 replicates per genotype per condition.

### Heat stress induces pronounced alterations in mRNA stability in Atdus2

Next, we aimed to understand the DHU-dependent molecular responses that make the *Atdus2* background less robust under abiotic stress. For this, we profiled *AtDUS2*-dependent transcripts and DHU abundance, as well as RNA-decay changes that occur in response to heat stress in Arabidopsis. Heat stress, rather than water withholding, was applied as it is both easier to apply consistently and quantitatively and it can be applied to seedlings in the RNA-decay experimental framework. In addition, our data indicated that *Atdus2* mutants showed diminished growth/survival at both the seedling and vegetative rosette stages under heat stress (**Figures 5D** and **S4D**). We monitored DHU abundance using HAMR/Modtect RCM calling of Illumina RNA-seq collected daily over a period of four days following implementation of heat stress (*N* = 8 per genotype and condition). An examination of these data revealed significantly increased mean DHU levels in Col-0, but not in *Atdus2*, in plants exposed to heat stress relative to their appropriate controls (**Figure 6A**, *p* = 0.048 and *p* = 0.88, respectively), suggesting that DHU is a standard mRNA-level stress response. As expected, we also observed a significant heat-stress associated increase in the other HAMR/Modtect identifiable RCMs in both Col-0 and *Atdus2* backgrounds (**Figure 6A**, *p* = 0.037 and *p* = 0.024, respectively). These data indicate a heat-dependent shift in DHU profiles that is dependent on *AtDUS2*.

**Figure 6.**
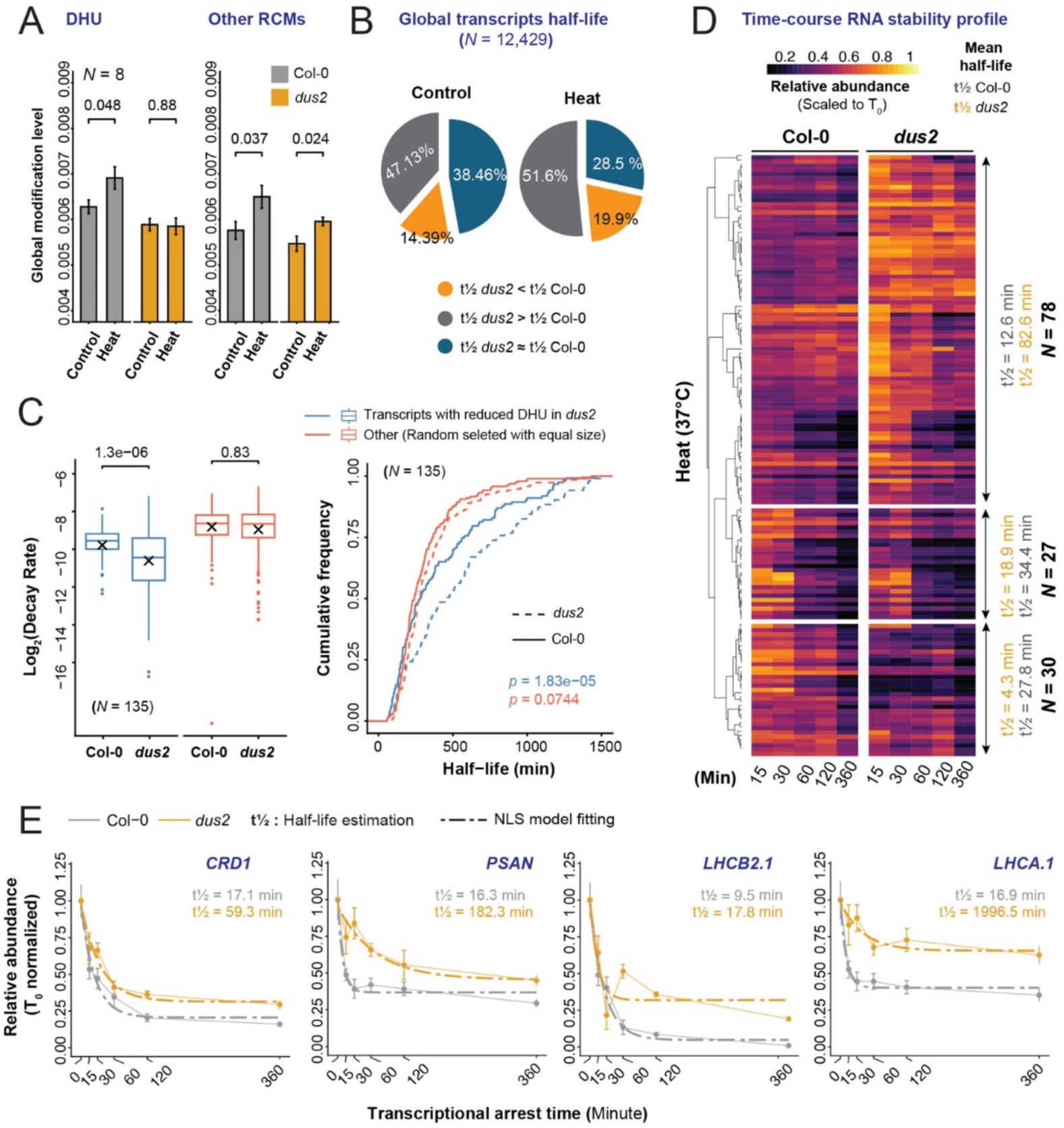
Comparison of abiotic stress-induced RNA stability between Col-0 and *dus2*. **(A)** A total of 12,450 transcripts with fitted half-life model were compared across two genotypes and two conditions, classified into six groups with respective proportions displayed using pie charts. Each group was classified based on log_2_FoldChange(half-life) based on exponential model fitting. **(B)** Left: Decay rate (*K*) based on exponential modeling on groups of transcripts with lower DHU level in *dus2* mutants under heat condition (blue boxplots) was compared with subset of randomly selected transcripts with equal sample size (red boxplots) using t-test (*p* < 0.05). Right: Cumulative frequency of estimated half-life of transcripts in Col-0 (solid line) and *dus2* (dashed line) background with lower DHU level in *dus2* mutants (blue lines) were compared with randomly selected transcripts (red lines) under heat stress. **(C)** Heatmap of 20 transcripts associated with differential DHU, transcript abundance, and protein abundance was plotted. Boxplots show GMUCT and RNA modification level of 16 transcripts with reduced protein in Col-0 relative to *dus2* (Wilcox test, *p* < 0.05). **(D)** Heatmap of 18 transcripts associated with differential DHU under both 22°C and 37°C conditions (left), protein class (photosystem I/II) were indicated with yellow and green box, Blue/grey colored boxes along the right indicate whether the Arabidopsis transcripts have a DHU modified ortholog in sorghum, and whether that transcript was assigned to the “blue” module or not (grey). **(E)** Box plot of DHU level and other RCM level in Col-0 (grey) and *dus2* mutants (orange) under the two conditions (t.test, *p* < 0.05). **(F)** Estimated RNA half-life of 15/18 transcripts with differentiated DHU level and degradation in Col-0 and *dus2*, facet by genotypes, with half-life estimated using exponential model and compared among four groups (Wilcox test, *p* < 0.05).

We then profiled the transcriptomic changes that occurred in these two backgrounds following heat stress. A total of 3,186 and 3,259 transcripts (**Table S14**) were differentially abundant in Col-0 and *Atdus2*, respectively, between control and heat stress conditions (*FDR* < 0.05, |log_2_FC| > 1; **Figure S9A and S9C**). As expected, the transcripts up-regulated in both backgrounds (**Figure S9B**, grey bars) were enriched for stress response-related terms, indicating the applied stress was sufficient to elicit a response However, terms enriched in the set of Col-0 specifically upregulated transcripts suggest that it is mounting a response to DNA damage, recombinational repair, and slowing down cell division, whereas the *Atdus2* mutant exhibited an increase in jasmonic acid related terms and a decrease in salicylic acid and photosynthesis-related terms (**Figure S9D**). To elucidate genotypic-specific stress response due to *Atdus2* loss-of-function, we profiled the unique transcripts that were differentially abundant in response to heat stress (**Figure S7C**). Particularly, these are subset of transcripts that exhibited a strong stress response in the Col-0 but were unaltered in the *Atdus2* background (**Figure S7C**, red boxes), suggesting that a functional DUS2 was necessary for their heat-associated response. We profiled the top three non-redundant KEGG and GO terms of transcripts either up or down-regulated in Col-0 in response to heat (**Figure S7D**). As expected, photosynthesis was the top enriched KEGG and GO term in the down-regulated transcripts in Col-0 in response to heat. In sum, these data suggest that Arabidopsis DUS2 is necessary for an appropriate DHU-mediated stress response, with photosynthesis-related traits and transcripts the primary target.

To further explore the mechanism driving the increased stress susceptibility observed in the *Atdus2* background, we next assessed if DHU-associated changes might influence RNA stability during heat stress. To assess RNA stability, seven-day-old seedlings in ½ MS liquid culture were acclimated at 37°C for 12 hours, and then treated with cordycepin, with samples collected over six hours as we did under control condition (**Figure S6B**). Global profiling of mRNA decay rates for the 12,450 model-fitted transcripts (**Table S12**) revealed that, in response to heat stress, a substantial proportion of mRNAs exhibited longer half-lives in *Atdus2* compared with Col-0 under both conditions (∼47% under control and 51.6% under heat (log₂FC(half-life ratio) > 0.1; **Figure 6B**). We than focused on transcripts with detectable DHU in Col-0 under heat stress. Using the HAMR/Modtect approach, we identified a total of 357 DHU-modified transcripts in Col-0 under heat, 213 of which showed reduced DHU in *dus2* (**Table S12**). Further selecting for transcripts with fitted decay models, we obtained 135 transcripts for comparison. For these transcripts, we observed significantly reduced decay rates (*K*) under heat in the *Atdus2* background (*p* = 1.3e-06, blue boxplots) that was not evident for non DHU-modified transcripts (red boxplots; **Figure 6C, left**, *p* = 0.88). This result is consistent with the cumulative frequency of half-lives, where transcripts associated with reduced DHU level in *Atdus2* exhibited significantly shorter half-lives in Col-0 than *dus2* (*p*= 1.83e-05, blue lines). In contrast, the half-lives of non-DHU modified transcripts remained unchanged between the two genetic backgrounds (**Figure 6C, right,** *p* = 0.0744, red lines).

A closer examination of these 135 transcripts using clustering heatmap uncovered three groups of transcripts with differentiated decay profiling (**Figure 6D**). More than half of transcripts (*N* = 78) exhibited substantially reduced half-lives in Col-0 relative to *Atdus2* in response to heat stress (**Figure 6D and Figure S8A**, average half-lives denoted for each cluster), whereas the remaining 57 transcripts exhibited increased half-lives in Col-0 under heat. In control settings, the DHU-modified transcripts with increased half-lives in the *Atdus2* background were enriched for the expected photosynthetic terms (*N* = 125, **Figure S8B**, top). Photosynthetic terms were also enriched for the pool of 78 DHU-modified transcripts with increased half-lives under heat in the *Atdus2*, but were also accompanied by abiotic stress terms (**Figure S8B,** bottom). Of the 78 DHU-modified transcripts that displayed a longer half-life in the *Atdus2* mutants relative to Col-0 under heat stress, 33 were associated with photosynthesis terms, including common photosystem I and II-associated transcripts, (eg., LHCA, LHCB, and PSB transcripts). Indeed, these transcripts had some of the largest half-life shifts, ranging from 2-100x longer (**Figure 6E**), indicating that DHU is necessary for their faster turnover in Col-0. In sum, these data suggest that the diminished performance under heat stress is likely due to impaired turnover of these photosynthesis-related transcripts and point to a novel post-transcriptional regulatory mechanism for these critical genes.

## Discussion

Drought and heat are common abiotic stressors that induce complex phenotypic and molecular responses that often lead to reduced photosynthesis, biomass, and seed set, thereby adversely affecting the overall productivity of agricultural crops. Sorghum’s adaptation to heat and drought makes its genetic and phenotypic diversity useful for revealing regulatory mechanisms that enable maximal growth under diverse field conditions. Although a variety of stress-responsive genes have been identified in sorghum and other stress resilient plant species (Pardo and VanBuren, 2021; Pardo et al., 2023), gaps remain, particularly in post-transcriptional regulatory events that have been shown to impact performance in other species.

To connect differences in plant performance in the field to specific transcriptional and post-transcriptional regulatory events, we performed a multi-omic comparison of six diverse sorghum accessions grown under water limiting and control conditions. We integrated molecular and physiological variation in a systems-level analysis to identify potential causal factors. As expected, physiological parameters and genes associated with photosynthesis, water movement, and stress-response regulation were all impacted to varying degrees in our panel (**Figure 1**). Interestingly, we also observed variation in one of the RCMs that we were able to be detected through standard Illumina RNA-sequencing approaches, DHU, that was strongly correlated with transcript abundance and plant performance in the six accessions (**Figure 2A**). Furthermore, our co-expression network analysis identified a dihydrouridine synthase (*SbDUS2*) transcript that was also associated with photosynthesis traits and DHU status of related transcripts. Importantly, the correlation between *SbDUS2* expression and the suite of transcripts with predicted DHU sites was observed in other public datasets and was not unique to our specific field conditions, growth parameters, or accessions. Thus, SbDUS2 and DHU appear to be linked with transcript groups associated with physiological traits.

While RCMs have been shown to influence or be associated with plant stress responses (Yu et al., 2021; Sharma et al., 2023; Cai et al., 2025), the identification of SbDUS2 associated with photosynthesis and stress-related transcripts was nevertheless a surprise. The DHU modification is mostly associated with tRNAs, and indeed the DUS protein family is conserved across the tree of life, including in eukaryotes, prokaryotes, and some archaea (Yu et al., 2011; Finet et al., 2022a; Finet et al., 2022b). Due to advances in detection capabilities, DHU has recently been described on mRNAs in humans and yeast (Dai et al., 2021; Finet et al., 2022b; Draycott et al., 2022), where it has been reported to influence splicing, translation, and fundamental biological processes such as meiosis (Finet et al., 2022b). Using computational algorithms that infer DHU status based on non-random errors introduced during the RT-PCR step of RNA-seq library preparation, we were also able to predict DHU-marked transcripts, and indeed our data suggest that these transcripts were enriched for a set of photosynthesis, metabolism, and other fundamental cellular processes. Whether this is due to sequence intrinsic features, functional conservation, or speculatively is an ancient organellar feature from before they were incorporated into the nuclear genome, is unclear and will likely guide future discoveries. The enrichment of predicted DHU on these highly abundant transcripts, may also be due to detection biases that scale with abundance. Monitoring DHU at the transcript level, or on lowly abundant transcripts, will likely only be possible with technical advances, such as ONT’s direct RNA-seq, and associated detection algorithms.

Given that this was the first detailed study of DHU in plants, and in particular on how DHU might contribute to plant growth and development, we sought to develop a clearer picture of how DHU is performing this function by examining Arabidopsis lines with the orthologous *Dus2* locus disrupted. Using LC-MS/MS, we confirmed that DHU levels were reduced in total RNA derived from *Atdus2* mutants relative to the Col-0 controls. Since we expected much of this DHU signal to be associated with the more abundant tRNA class, we also specifically examined mRNA-associated DHU levels using both HAMR/Modtect on Illumina RNA-seq and by comparing Nanopore DRS between WT (Col-0) and *Atdus2* mutants. We observed a DUS2-dependent decrease in DHU levels in mRNA using these two complementary approaches, suggesting that AtDUS2, like other eukaryotic DUS2 enzymes, modifies mRNAs in addition to other RNA classes. With evidence supporting a DUS2-dependent reduction in mRNA-associated DHU, we then assessed the impact that diminished DHU abundance had on plant performance. A very real concern is that the loss of DUS2 might have pleotropic effects on plant performance simply due to its potential role in modifying tRNA, thereby diminishing translation of all protein classes. Importantly, untargeted protein mass spectrometry did not reveal a broad reduction in protein abundance **(Figure S6**) but instead revealed a very specific reduction in proteins associated with photosynthesis and metabolism, similar to the class of mRNAs associated with DHU modification. Whether AtDUS2 doesn’t play a prominent role in the addition of DHU to tRNAs, or if its absence is compensated by one of the other four DUS enzymes in Arabidopsis, remains to be tested.

An examination of the transcriptomic and proteomic changes in the *Atdus2* background provided clues as to what might give rise to the reduced germination and delayed development under control conditions. We predominantly detected a reduction in photosynthesis-related transcripts in the *Atdus2* background. Dysregulation of these transcripts could certainly explain the delayed, slower growth that we observed in both Arabidopsis and in sorghum with reduced *DUS2* levels (e.g., the “A” accession), and under stress or control conditions, as plants with reduced photosynthesis struggle to efficiently capture and convert light energy into glucose. Interestingly, glucose, alongside other carbohydrates, were strongly correlated with the photosynthesis-associated *DUS2* co-expression modules **(Figure 2A**), again highlighting a model where DUS2, and DHU, are potentially critical for efficient photosynthesis.

The observed impact of *AtDus2-*dependent DHU on photosynthesis were pronounced under abiotic stress conditions in Arabidopsis, with reduced *AtDUS2* levels associated with decreased plant performance. This raises the question of how a modification that appears to be negatively associated with transcript abundance could be beneficial for plant performance under stress? We propose a model, supported by RNA decay data in both Col-0 and *Atdus2* mutants under control and heat conditions, whereby DHU is likely necessary for appropriate mRNA turnover under stress conditions (**Figure 7**). The observation that DHU-associated photosynthesis transcripts exhibited substantially increased half-lives (up to 100-fold) in the Atdus2 background provides support for this model of DHU-dependent transcript destabilization. In addition, prior work examining HAMR detectable RCMs in Arabidopsis found that this group of modifications, which included DHU, were heavily enriched on actively degrading transcripts under control conditions (Vandivier et al., 2015). Higher turnover, reduced steady state levels, and structural perturbations of photosynthesis and metabolism related transcripts have been observed under stress conditions in a number of plant systems (Li et al., 2022; Notarnicola et al., 2023) with these changes proposed as a rapid means of clearing molecules that are accumulating damage caused by reactive oxygen species (ROS; Park et al., 2012). While plants attempt to limit photosynthesis during stress, as it is one of the main ROS producing mechanisms in the cell, inappropriately elevated photosynthesis can lead to cell death-inducing ROS levels. We propose that DHU is an RCM that marks mRNAs for degradation, thereby might helping to clear them more rapidly and facilitate a proper response to environmental perturbations. Excitingly, there is a clear connection between the cytoplasmic mRNA decay machinery and stress tolerance in Arabidopsis (Merret et al., 2013) and DHU, or a sufficient accumulation of DHU marks, may target mRNAs towards this pathway. We propose that Arabidopsis plants lacking DUS2, or that decrease DUS2 levels or activity in response to stress conditions, would then be unable to clear these damaged mRNA molecules through cytoplasmic decay pathways, potentially increasing cell death and diminishing plant performance.

**Figure 7.**
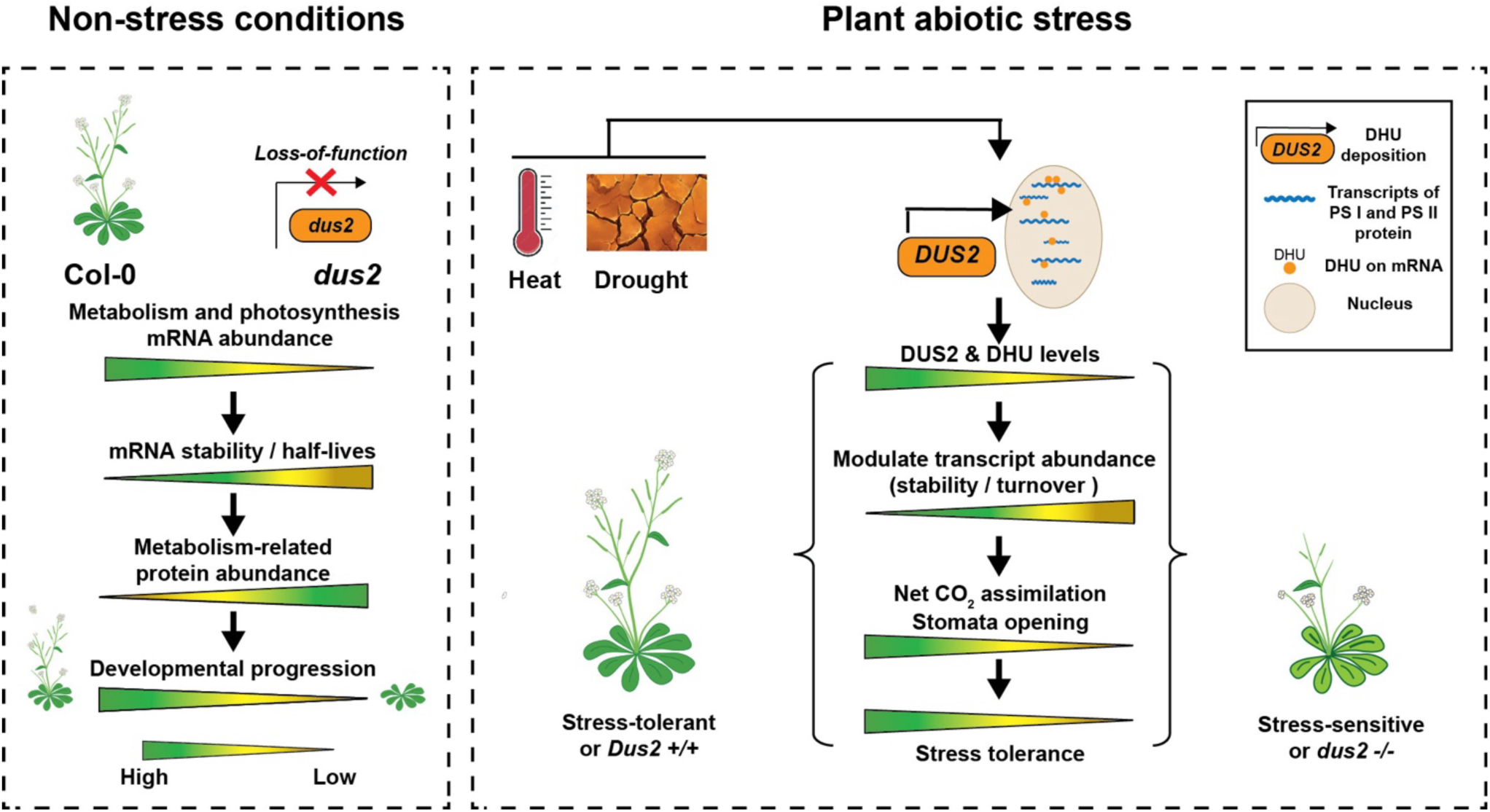
Proposed model explains DHU impacts on plant growth and stress response. Based on our findings in *Arabidopsis*, reduced DHU levels appear to impact the abundance of a subset of DHU-modified transcripts in Col-0. Loss of DUS2 shifts the stability and half-lives of mRNAs encoding proteins involved in photosynthesis and metabolism, which in turn leads to impaired germination and delayed growth in the mutant background. Growth defects are exacerbated when plants are grown under abiotic stress conditions, due to a proposed inappropriate turnover of photosynthesis-associated transcripts. Counterintuitively, increases in these unmodified photosynthesis-related transcripts in the Atdus2 mutant leads to reduced photosynthesis parameters, likely due to a buildup of ROS and eventual cell death. Thus, DHU is necessary to reduce mRNA steady state levels, or to target mRNAs for degradation, helping to manage the pool of photosynthesis-related proteins and ultimately allowing for better stress tolerance.

In summary, our comparative, multi-species approach facilitated the identification of an RCM that appears to be critical for plant performance under stress. Indeed, this RCM influences abiotic stress responses in Arabidopsis by acting on many of the photosynthesis and metabolism-related mRNAs. These data highlight the complex transcriptional and post-transcriptional regulatory landscape influencing plant stress responses and offer promising insights as to how they might be used to select more stress-resilient plants in the future.

## Supporting information

All supplementary tables

## Data availability

Transcriptome sequencing reads have been deposited in the National Center for Biotechnology Information (NCBI) BioProject database under the accession number: PRJNA1119650. Chromatograms of 5,6-dihydrouridine Mass spectrometry data was deposited under massIVE online repository (ftp://massive.ucsd.edu/v09/MSV000097205/). Computational pipelines used for data analysis were deposited under the GitHub page (https://github.com/Leon-Yu0320/Sorghum_Range28). The R markdown files (R code) for data visualization in main figures and supplementary figures were deposited in Rpubs workspace (https://rpubs.com/LeonYu/Main-Figures and https://rpubs.com/LeonYu/Supp-Figures).

## Acknowledgments

We would like to thank the members of the epitranscriptomic working group within the Nelson, Pauli, Schroeder, and Gregory labs for their insightful comments. We would like to thank Dr. Cliff Weil for insightful comments about the SAP, Dr. Tony Schilmiller and the MSU Mass Spectrometry and Metabolomics Core for assistance in metabolomics measurements and analysis. Many thanks to Dr. Alyssa Kearly for her assistance in technical proofing of the manuscript. This work was supported by NSF IOS PGRP 2023310 (to ADLN, EHL, DP, and BDG), NSF IOS PGRP 1849708 (to BDG), NSF MCB 2427729 (to BDG), NSF MCB 2102120 (to ADLN and DP), Department of Energy (DOE) DE-AR0001101 (to DP), DE-SC0023305 (to DP and AE), DE-SC0020401 (to AE and DP), Cotton Incorporated 18-384, 20720, and 21-830 (to DP), NSF DBI-2019674 (to DP and ADLN), and NIH R35GM153298 (to AIS).

## Author contributions

LY, GM, AEA, EHL, BDG, ADLN, and DP developed the experiment. GM, SC, BR, and DP designed the field experiment and collected field-related data. ACND and EKB extracted RNA and performed other sorghum-related molecular experiments. AE provided accessions from the SAP for this study. LY, ApS, XZ, and EKB worked with the Arabidopsis DUS2 mutants. DRG generated and contributed to the analysis of GMUCT data. GM and TO generated the metabolomics data. HA and GB generated the biochemical data related to oxidative stress status. LY, RH, and DS helped and provided expertise in acquiring physiological measurements for the Arabidopsis DUS2 mutants. HF and AlS performed mass-spec experiments on the DUS2 mutants. LY analyzed all RNA-seq and developed co-expression networks and data integration. LY, KP, and SJS analyzed epitranscriptomic data. LY, BDG, ADLN, and DP wrote the manuscript.

## Supplementary Figures

**Figure S1. Genomic composition of variants of six sorghum accessions:** 8,725,449 variants (SNPs and INDELs) derived from variants calling (GATK4, reference genome: **McCormick et al., 2018)** were annotated based on genomic classifications using VEP tool, the unique variants of each accession were calculated across upstream, downstream, intron, and splicing regions.

**Figure S2. Maximum likelihood phylogeny of plant and budding yeast DUS proteins**

A maximum likelihood phylogeny was inferred from a protein alignment of three DUS genes identified in Arabidopsis (AT), Sorghum (Sobic), Oryza sativa (rice, Os), Physcomitreum patens (Pp), and Saccharomyces cerevisiae (Sc) through reciprocal best BLASTp. Values at each node represent bootstrap support (out of 100 replicates).

**Figure S3. Features of co-expression modules_in sorghum field data**

**(A)** A child network (*N* = 48) was displayed by including two DUS genes (orange box) and intersected genes derived from 326 DHU-modified transcripts and photosynthesis and metabolic pathway enriched genes (*N* = 96). Transcripts with a correlation between DHU level and transcript abundance were marked in light blue (*N* = 29), with orange dashed lines connecting DUS-derived connections and light red dashed lines indicating known protein-protein interactions from the STRING database. **(B)** Heat map of 29 transcripts that displayed a negative correlation between DHU and transcript abundance. Correlation between DHU and transcript abundance is shown across the top of the heatmap, with the correlation between DUS2/3 and DHU modified transcript abundance shown along the bottom. **(C)** Examination of expression for these 29 transcripts and *SbDUS2*/*SbDUS3* in sorghum tissues or developmental stages using publicly available tissue expression atlas data from McCormick et al., 2018. Correlation between DHU and transcript abundance is shown across the top of the heatmap, with the correlation between DUS2/3 and DHU modified transcript abundance shown along the bottom. **(D)** Normalized transcript abundance (TPM) values of DUS2 and DUS2 in the sorghum public tissue atlas**. (E)** Pearson correlation between SbDUS2/3 transcript abundance relative to the PC1 score of the PCA derived from DHU-level of 326 DHU-modified transcripts in “blue” module. **(F)** The TPM level of *SbDUS1* transcripts in sorghum drought experiment.

**Figure S4. Characterization of *Atdus2* mutants**

**(A)** Illumina-mapped reads and DRS reads from DUS2 mutants and Col-0 background were visualized using the IGV tool to confirm mutant samples contain homozygous T-DNA insertion that disrupts the expression of full-length *DUS2*. **(B)** Illustration of water holding experiment to maintain certain soil water content over drought treatment. **(C)** Drought stress on 4-week Arabidopsis plants at 70% SWC or 25% SWC over time using Col-0 and two independent *dus2* mutant lines (CS857388 and SALK_079129C). **(D)** Heat stress on 7-day-old Arabidopsis seedlings grown on MS plates under control condition and prolonged heat stress 37°C (48 hours) using Col-0 and two independent *dus2* mutant lines (CS857388 and SALK_079129C). **(E)** Bud initiation rate of Col-0 and *Atdus2* plants: Percent of plants with flower buds visible plotted over time are recorded. **(F)** The plant architecture of Col-0 and *Atdus2* plants at about 50 days of growth.

**Figure S5. DHU deposition characteristics in sorghum and Arabidopsis**

**(A)** Frequencies of all 3-nt sequence combinations adjacent to DHU-modified sites (red) in sorghum and *Arabidopsis* are summarized. U-tract sites are defined as those containing at least one adjacent U near the DHU position. **(B)** Metaplots showing the distribution of DHU-modified sites in sorghum and *Arabidopsis* across the 5′UTR (±250 bp), 3′UTR (±250 bp), and full transcript bodies.

**Figure S6. Molecular features of DHU-modified transcripts in Arabidopsis**

**(A)** The volcano plot shows up and downregulated transcripts in the mutant background. **(B)** The volcano plot shows proteins associated with increased and decreased abundance in the mutant background. **(C)** Log_2_FC (*dus2*/Col-0) of 275 transcripts and proteins were plotted to show correlation between transcripts and proteins, the DHU modified transcripts were marked in “dark yellow” box. **(D)** Log_2_FC (*dus2*/Col-0) of DHU and protein abundance were plotted to show correlation between DHU and protein level. **(E)** Protein abundance (based on MS) and transcript abundance (RNA-seq) derived from paired samples. Only genes observed in RNA-seq, MS, and for which DHU sites can be inferred, are shown. Red dashed box highlights transcripts with increased abundance in *Atdus2*. **(F)** Density plot of exon-exon junction footprint of GMUCT data.

**Figure S7. Heat-induced transcriptome response and RNA stability assay**

**(A)** Cumulative frequency plot presents the distribution of the log_2_FC(*dus2*/Col-0) of transcripts with lower DHU level in *dus2* background relative to other transcripts under control and heat conditions. *p*-value was obtained using a two-sided Wilcoxon rank sum test. **(B)** 7-day-old seedling of two Arabidopsis genotypes were grown on ½ MS media and transformed to liquid media for 12-hour incubation under 22°C and 37°C conditions. Samples were collected at various time points (0, 15, 30, 60, 120, 360 min) after cordycepin treatment for RNA extraction and sequencing **(F)** Upset plot displaying the number of all up- or down-regulated heat-induced transcripts in Col-0 and *dus2* genotypes, with red boxes highlight transcript present uniquely in one genotype at certain condition **(G)** KEGG and GO term enrichment (-log_10_ *FDR*) of Col-0-specific DATs (from panel G). Only the top three terms are shown for up or down-regulated transcripts, with the number of enriched transcripts relative to the total transcripts shown in, or adjacent, to each box.

**Figure S8. Heat-induced transcriptome response in Col-0 and *Atdus2* plants**

**(A)** The volcano plot shows up and downregulated transcripts (heat / control) in the wildtype plants. **(B)** The bar plots show -log_10_(*FDR*) and the number in red display fold enrichment of each 10 enriched terms involved in biological processes. **(C)** The volcano plot shows up and downregulated transcripts (heat / control) in mutant plants. **(D)** The bar plots show -log_10_(*FDR*) and the number in red display fold enrichment of each 10 enriched terms involved in biological processes.

**Figure S9. RNA stability and GO enrichment of subset transcripts under two conditions**

(A) NLS modeling of subset transcripts derived from 216 (135) transcripts with reduced DHU level in *dus2* background under control (heat) conditions. Each three groups under control and heat conditions were divided based on clustering of decay profile of transcripts. (B) GO enrichment of two larger group of transcripts with faster decay in Col-0 than *dus2* (*N* = 125 under control conditions and *N* = 78 under heat conditions).

